# Distinct Contribution of DNA Methylation and Histone Acetylation to the Genomic Occupancy of Transcription Factors

**DOI:** 10.1101/670307

**Authors:** Martin Cusack, Hamish W. King, Paolo Spingardi, Benedikt M. Kessler, Robert J. Klose, Skirmantas Kriaucionis

## Abstract

Epigenetic modifications on chromatin play important roles in regulating gene expression. While chromatin states are often governed by multi-layered structure, how individual pathways contribute to gene expression remains poorly understood. For example, DNA methylation is known to regulate transcription factor binding but also to recruit methyl-CpG binding proteins that affect chromatin structure through the activity of histone deacetylase complexes (HDACs). Both of these mechanisms can potentially affect gene expression, but the importance of each, and whether these activities are integrated to achieve appropriate gene regulation, remains largely unknown. To address this important question, we measured gene expression, chromatin accessibility, and transcription factor occupancy in wild-type or DNA methylation-deficient mouse embryonic stem cells following HDAC inhibition. Interestingly, we observe widespread increases in chromatin accessibility at repeat elements when HDACs are inhibited, and this is magnified when cells also lack DNA methylation. A subset of these elements have elevated binding of the YY1 and GABPA transcription factors and increased expression. The pronounced additive effect of HDAC inhibition in DNA methylation deficient cells demonstrate that DNA methylation and histone deacetylation act largely independently to suppress transcription factor binding and gene expression.

## Introduction

The most abundant DNA modification in mammals is 5-methylcytosine (5mC) which is found predominantly within CpG dinucleotides (Lister et al., 2009; Stadler et al., 2011). DNA methylation is deposited and maintained through the concerted activity of three essential DNA methyltransferases (DNMT1, DNMT3A, DNMT3B) (Goll and Bestor, 2005) and plays important roles in transcriptional repression. For example, DNA methylation is particularly important for the regulation of imprinted genes (Kaneda et al., 2004), some tissue-restricted genes (De Smet et al., 1999; Maatouk et al., 2006; Borgel et al., 2010), evolutionarily young repeat elements (Davis et al., 1989; Walsh et al., 1998), genes on the inactive X chromosome (Beard et al., 1995; Hansen et al., 1996) and certain tumour suppressor genes in cancer cells (Toyota et al., 1999; Rhee et al., 2002; Robert et al., 2003).

Although 60-80% of CpGs in the genome are methylated, DNA methylation is absent or reduced in regions bound by transcription factors such as CpG islands, gene promoters and distal regulatory elements (Stadler et al., 2011; Hon et al., 2013; Ziller et al., 2013). This observation led to the proposal that DNA methylation may limit transcription factor occupancy. In support of this model, *in vitro* experiments have revealed that 5mC can affect the affinity of most DNA-binding proteins that contain CpG dinucleotides in their binding motif (Hu et al., 2013; Kribelbauer et al., 2017; Spruijt et al., 2013; Yin et al., 2017). Furthermore, certain transcription factors (TFs), including NRF1 and CTCF, have been shown to have altered binding profiles in cells where DNA methylation is reduced or absent (Domcke et al., 2015; Maurano et al., 2015; Yin et al., 2017).

In addition to directly altering the binding of transcription factors, methylated CpG is also recognised by methyl binding domain-containing (MBD) proteins (Hendrich and Bird, 1998). MBD proteins interact with histone deacetylase (HDAC) enzymes and promote the formation of transcriptionally repressive chromatin environments (Kokura et al., 2001; Nan et al., 1998; Yoon et al., 2003; Zhang et al., 1999) via the removal of acetyl groups from lysine residues in histones. Deacetylation is thought to contribute to gene repression through a number of mechanisms. Firstly, the absence of acetyl groups on lysines means they can be methylated and recruit methylysine-binding proteins, some of which counteract transcription (Bannister et al., 2001; Lachner et al., 2001). Secondly, deacetylated histones contribute to chromatin compaction through strengthening histone tail-DNA interactions (Hong et al., 1993; Lee et al., 1993; Anderson et al., 2001; Wang and Hayes, 2008). Thirdly, the absence of acetylation on lysine residues, limits the binding of bromodomain-containing transcriptional activators, many of which recognise acetylated histone tails (Dhalluin et al., 1999).

Despite the active recruitment of HDACs to methylated CpGs, the extent to which DNA methylation relies on HDAC activity to affect gene expression is still poorly understood. HDAC inhibition, or disruption of the interaction between MBD proteins and the HDAC-containing complexes, have been shown to alleviate transcriptional silencing mediated by DNA methylation in reporter gene assays (Jones et al., 1998; Lyst et al., 2013; Nan et al., 1998; Ng et al., 1999). Nevertheless, at other endogenous genes, inhibiting HDACs did not recapitulate the effects on gene expression that manifest when DNA methylation is removed (Brunmeir et al., 2010; Cameron et al., 1999; Coffee et al., 1999; Reichmann et al., 2012). Therefore, while it is clear that DNA methylation is capable of altering TF-DNA interactions and recruiting HDACs, how these distinct mechanisms contribute to DNA methylation-dependent repression at the genome-scale remains unknown.

Here we define the effects of either loss of DNA methylation or histone deacetylase activity on transcription factor occupancy and gene expression. To achieve this we profiled chromatin accessibility, transcription factor occupancy, and gene expression in wild-type (WT) and DNA methylation-deficient mouse embryonic stem cells in the presence or absence of HDAC inhibition. We anticipated that HDAC inhibition would mirror the effects of DNA methylation loss at genomic loci for which HDAC activity is required for DNA methylation-mediated repression. However, surprisingly, we discover that DNA methylation and HDACs acts largely as independent mechanisms to restrict TF occupancy in mouse ES cells. Interestingly, the relocation of TFs and the accompanying changes in accessibility caused by loss of DNA methylation and HDAC inhibition only rarely affected the activity of proximal genes. In contrast, YY1 and GABPA occupancy at LINE and LTR repeats correlated with elevated expression from these retrotransposons.

## Results

### Disruption of HDAC activity and DNA methylation in mouse embryonic stem cells

To determine the contribution of DNA methylation and the activity of HDAC enzymes on transcriptional regulation, we treated wild-type (J1) or *Dnmt3a*/*Dnmt3b*/*Dnmt1* triple knockout (DNMT.TKO) mouse embryonic stem cells (Tsumura et al., 2006) with HDAC inhibitor (Figure 1A). The chosen dose of trichostatin A (TSA) had minimal effects on the morphology of mESCs, but yielded a robust increase in global histone acetylation as detected by immunoblotting (Figure 1B). There were no significant changes in global 5mC levels between TSA- and DMSO(solvent)-treated cells (Figure S1A).

**Figure 1:**
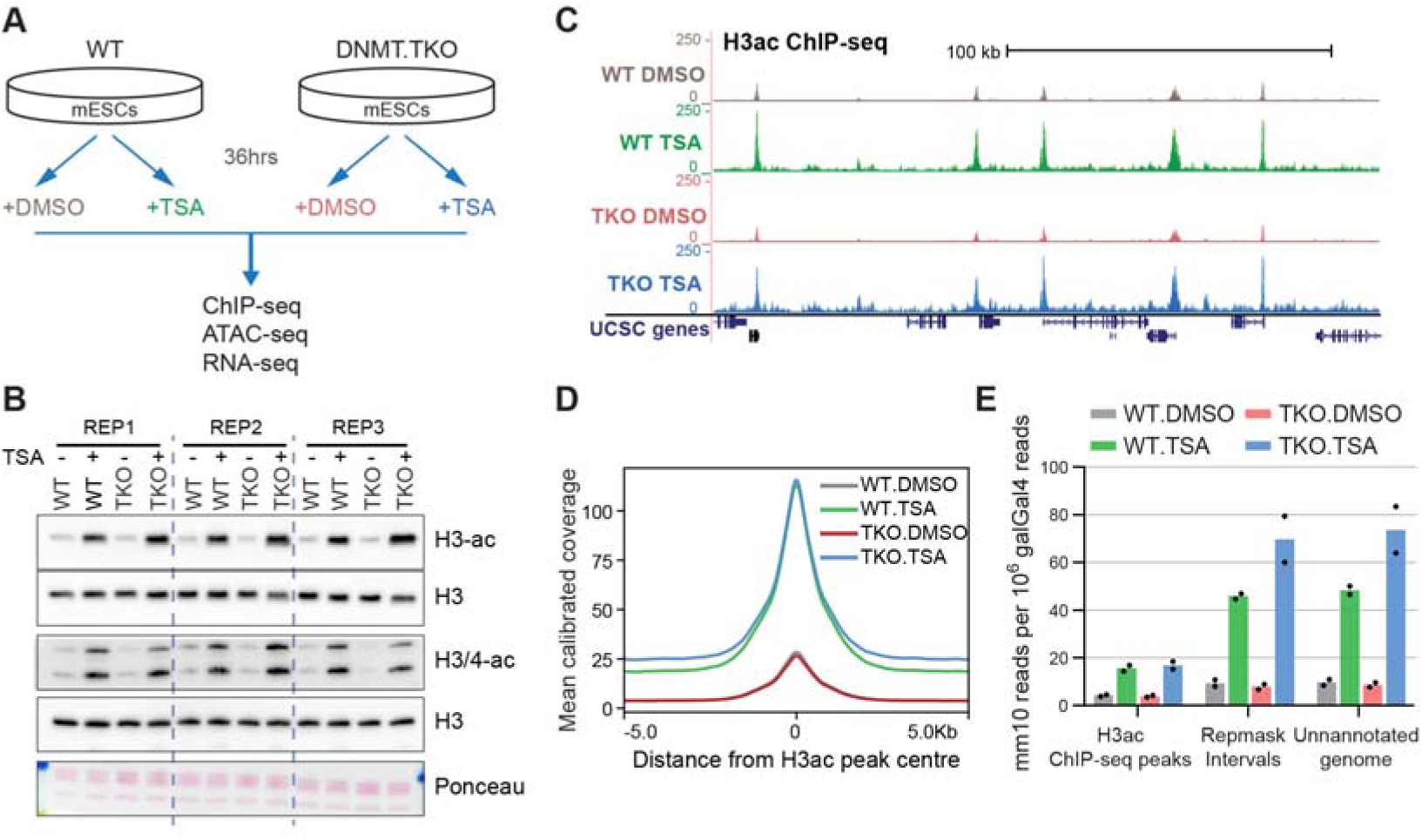
Disruption of HDAC activity and DNA methylation in mouse embryonic stem cells. **A.** Schematic of the experimental approach used in this study. **B.** Immunoblot analysis of global histone H3 and H4 acetylation levels after 36 h TSA treatment in mESCs. Samples were derived from three biological replicate experiments. **C.** Representative UCSC genome browser snapshot showing H3ac ChIP-seq read coverage calibrated to the spike-in galGal4 genome. Coverage graphs were generated as described in Materials and Methods after merging alignments from two replicate samples. Mm10 coordinates: chr3:94,876,915-95,075,953. **D.** Metaplot showing the average calibrated H3ac ChIP-seq signal from DMSO or TSA-treated wild-type and DNMT.TKO cells in 10 kb regions surrounding the centre of ChIP-seq peaks (n = 28608). **E.** Distribution of H3ac ChIP-seq reads within genomic intervals separated into three mutually exclusive categories. For every sample, the number of mm10 reads overlapping each category was divided by the total number of reads mapping to the galGal4 genome. Data are represented as mean; points indicate the values for two biological replicates.

To examine histone acetylation in more detail, we performed native calibrated ChIP-seq for acetylated histone H3 (H3ac). TSA treatment led to a global increase in H3ac in wild-type and DNMT.TKO cells both at previously hyper-acetylated sites (ChIP-seq peaks called using MACS2 (Zhang et al., 2008)) and the remaining genomic space (Figure 1C, 1D and 1E). Interestingly, the increase in histone acetylation at repetitive regions and unannotated genomic loci following HDAC inhibition was greater in DNMT.TKO compared to wild-type cells (Figures 1E). This observation coincides with high DNA methylation levels in the aforementioned regions in WT cells, whereas hyper-acetylated sites contain low median DNA methylation (Figure S1C). This additive effect of DNA methylation loss and HDAC inhibition on histone acetylation in these regions could be explained by enhanced recruitment of histone acetyltransferases (HATs), potentiated by DNA binding proteins with specificity to unmodified CpGs (e.g. CxxC-domain proteins). Alternatively, the lack of DNA methylation may disrupt the recruitment of HDACs by MBD-domain proteins, resulting in a lower capacity for compensatory mechanisms to deacetylate histones or prevent histone acetylation.

In summary, HDAC inhibition using TSA induces histone acetylation broadly across the genome with no detectable change in DNA methylation, enabling us to evaluate the extent and the mode by which these two mechanisms contribute to chromatin function.

### DNA methylation and HDAC activity have distinct contributions to the chromatin accessibility landscape

To compare the impact of DNA methylation and histone acetylation on chromatin function we profiled chromatin accessibility in wild-type and DNMT.TKO cells in the absence or presence of TSA using the Assay for Transposase-Accessible Chromatin followed by high throughput sequencing (ATAC-seq) (Figure 1A).

From these data, we first defined a set of 83395 transposase hypersensitive sites (THS, also referred to as ATAC-seq peaks) that were called as significantly accessible in at least one experimental condition (FDR adjusted p-value below 10^−60^ using the DANPOS peak-calling algorithm (Chen et al., 2013). A principle component analysis that used ATAC-seq read counts at each THS interval clustered biological replicate samples together, demonstrating that treatment and genotype induced reproducible changes in accessibility (Figure S2A).

We first focused on comparing DNMT.TKO to WT DMSO-treated cells in order to characterise the chromatin accessibility changes that occur following the loss of 5mC. We identified 6427 THS regions that significantly gained accessibility and 7135 that significantly lost accessibility in the absence of DNA methylation (FDR < 0.01 and fold change > 1.5; Figure 2A - left). Differential THS regions account for ∼16% of all sites that were detected as being accessible in wild-type or DNMT.TKO mES cells, demonstrating that DNA methylation has an important role in maintaining accessibility patterns. However, complete loss of DNA methylation in mouse ES cells does not produce widespread alterations, in agreement with similar results obtained using DNAse-seq (Domcke et al., 2015).

**Figure 2:**
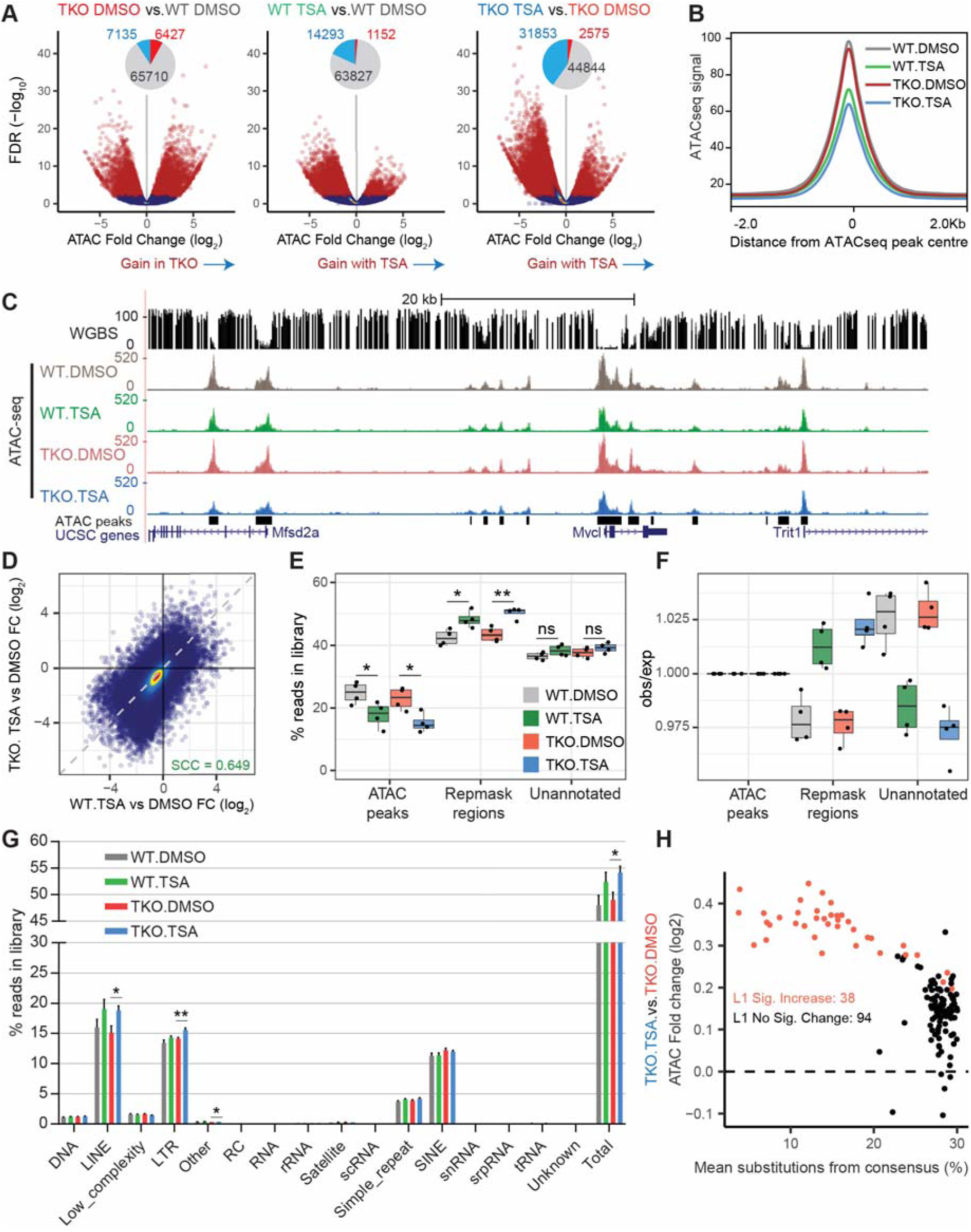
DNA methylation and HDAC activity have distinct contributions to the chromatin accessibility landscape. See also Figure S2. **A.** Volcano plots representing the false discovery rate (FDR) and fold change values obtained through pairwise differential analyses of ATAC-seq signal at 83395 THS. Regions with significantly differential accessibility (fold change > 1.5 and FDR < 0.01) are shown in red on the scatter plots and their numbers are summarised in the form of pie charts (light blue = significant decrease; red = significant increase). **B.** Metaplot showing the average ATAC-seq signal from DMSO- or TSA-treated wild-type and DNMT.TKO cells in 4 kb regions surrounding the centre of ATAC-seq peaks (n = 83395). **C.** Representative UCSC genome browser snapshot showing CpG methylation levels (Habibi et al., 2013) and ATAC-seq read coverage. THS intervals and genes are annotated. ATAC-seq coverage graphs were generated after merging alignments from four replicate samples. Mm10 coordinates: chr4:122,948,721-123,029,460. **D.** Scatterplot comparing the fold change in ATAC-seq signal following TSA treatment in DNMT.TKO versus wild-type cells. The dashed line has a slope of 1 and intercept of 0. Fold change values were obtained through differential analysis of the read coverage at 83395 ATAC-seq peaks. SCC = Spearman Correlation Coefficient. **E.** Boxplots summarising the percentage of reads from each ATAC-seq library that map to intervals split into three mutually exclusive categories. Two-tailed Student t-tests were used to determine significant differences:* p-value < 0.05; ** p-value < 0.01; *** p-value < 0.001; *ns* = non-significant (p-value > 0.05). **F.** Boxplots summarising the distribution of reads from each ATAC-seq library that map to intervals split into three mutually exclusive categories, relative to the distribution expected by chance i.e. if non-THS reads were shuffled randomly within the genomic space outside of THSs. For each sample, values shown in (E) are divided by those in Figure S2E. **G.** Distribution of ATAC-seq reads across different classes of repetitive elements. The number of reads that overlap each category was plotted as a percentage of the total library size. Data are represented as mean + SD. Two-tailed Student t-tests were used to determine significant differences between TSA and DMSO treated samples: * p-value < 0.05; ** p-value < 0.01; *** p-value < 0.001; non-significant differences are not indicated. **H.** Scatterplot comparing for each LINE-1 repeat subtype, the fold change in ATAC-seq signal to the average substitution rate (base mismatches relative to the consensus sequence in parts per 100). See also Table S3.

To assess the fraction of differential accessibility events that occur as a direct consequence of a loss in DNA methylation, we made use of public whole genome bisulfite sequencing (WGBS) data generated from E14 mESCs grown under the same growth conditions (Habibi et al., 2013). Similar to previous observations (Domcke et al., 2015), we found that gains in accessibility in DNMT.TKO cells frequently occur at regions that are methylated in WT cells (Figure S2B, top right quadrant). Specifically, 56% of DNMT.TKO loci that significantly gain in accessibility have a mean methylation value above 60% in WT cells; this percentage rises to 89% for the top 500 most differentially accessible regions (Figure S2B). These represent loci at which DNA methylation is likely to play a role in restraining the formation of a Tn5 accessible state, potentially through interference with transcription factor binding. On the other hand, the relationship between accessibility reduction in DNMT.TKO cells and DNA methylation is less straightforward (Figure S2B). Only 2.8% of the regions displaying significantly reduced accessibility in DNMT.TKO cells have high levels of methylation in wild-type cells (≥ 60% methylation), making it more difficult to attribute these differences in chromatin accessibility to changes in methylation.

Screening a collection of 264 known vertebrate transcription factor motifs (HOMER database, Heinz et al., 2010) for enrichment at DNMT.TKO-specific THSs, we identified the NRF1 motif as being the most enriched, consistent with the findings of Domcke et al., 2015 (Figure S2C and Table S5). Similar to their results, sequence motifs recognised by the NFY complex, ETS and basic Helix-Loop-Helix (bHLH) transcription factors were also enriched in DNMT.TKO THSs. *In vitro* affinity measurements and locus-specific ChIP evidence suggest that the binding of these transcription factors to their cognate recognition sites is sensitive to DNA methylation (Hu et al., 2017; Yin et al., 2017), supporting the idea that accessibility gains occur in the absence of DNA methylation due to unimpeded binding of these proteins.

Recent data from *in vitro* affinity assays, have revealed that certain TFs preferentially bind methylated versions of their cognate binding sequences (Kribelbauer et al., 2017; Mann et al., 2013; Rishi et al., 2010; Yin et al., 2017). If DNA methylation were required for binding of these TFs *in vivo*, the removal of 5mC would be expected to cause a loss of occupancy and thus reduced accessibility. Within 18% of THS, which are methylated (≥ 60% methylation) in WT cells, only a minority of sites (9.9%) show significantly decreased accessibility in DNMT.TKO cells (Figure S2B), implying that DNA methylation plays a limited role in facilitating the formation of accessible sites in ES cells.

We next examined the effects of HDAC inhibition on the chromatin accessibility landscape of wild-type and DNMT.TKO mES cells. To our surprise, few loci displayed significant accessibility gains in TSA-versus DMSO-treated cells, relative to the numbers identified when comparing DNMT.TKO to WT. In fact, we observed that the vast majority of regions display reduced accessibility in cells upon HDAC inhibition, in both WT and DNMT.TKO cells (Figure 2A). The effect was more pronounced in cells depleted of DNA methylation, in which 38% of all THSs showed a significant reduction in ATAC-seq signal when treated with TSA. Congruently, ATAC-seq peaks in TSA-treated cells are appreciably smaller, as seen when aggregating the signal across all THSs or at individual sites in the genome (Figures 2B and 2C). We further noted that genome accessibility changes (signal fold change at THS peaks) were similar between WT and DNMT.TKO cell lines (Figure 2D, Spearman Correlation Coefficient = 0.649), demonstrating that the main difference in the response to HDAC inhibition in either the presence or absence of DNA methylation is the magnitude of the changes.

Reduced signal at THS intervals in TSA-treated samples indicates that DNA fragments originating from these regions are underrepresented as a fraction of the entire ATAC-seq library, relative to their fraction in untreated samples. Concurrently, a larger proportion of fragments mapping to the inaccessible compartment of the genome (i.e. the background) are sequenced. We envision two possible explanations for this observed reduction of ATAC-seq signal at THS intervals.

The first possibility is that, upon TSA treatment, chromatin accessibility is reduced at THSs meaning that fewer reads would be obtained from within these intervals. As a consequence of sequencing a fixed number of DNA fragments, we would thus observe a relative increase in accessibility signal from the genomic compartment outside THSs (the ‘inaccessible’ compartment). Importantly, according to this hypothesis, the additional reads sequenced from the inaccessible compartment would be distributed evenly throughout the remaining genomic space. The second possibility is that TSA treatment results in elevated accessibility in genomic regions outside of THSs. The relative reduction in the ATAC-seq signal at THSs then becomes a secondary consequence of elevated accessibility outside THSs. This scenario would be supported if we were to identify an enrichment in accessibility at particular subdivisions of the inaccessible compartment. These two mechanisms are not mutually exclusive and both could contribute to the shift in read distribution observed in TSA-treated samples.

To explore the above-mentioned interpretations, we calculated the fraction of reads in the ATAC-seq libraries that overlapped with three categories of genomic regions: transposase hyper-accessible regions (the ATAC-seq peak set defined above, 83395 intervals covering 55 Mb, 2.03% of the mm10 genome); genomic intervals consisting of repetitive elements (RepMask intervals, excluding sequences that overlap THSs, 1.19 Gb, 43.39% of mm10 genome) and the remaining genomic space (unannotated regions, 54.58%) (Figure 2E). This analysis shows that gains in accessibility in TSA-treated samples occur preferentially at repetitive elements compared to unannotated regions (Figure 2E), supporting a contribution from the second scenario outlined above. The increase in coverage observed at repeats is larger than that obtained if the same number of non-THS sequencing reads are randomly shuffled into the combined “RepMask” and unannotated genomic compartments (Figures 2F, S2D and S2E). Furthermore, the distinct accessibility changes observed at repeat regions are unlikely to be solely a consequence of differences in the induction of histone acetylation as the H3ac levels in both compartments are equivalent following HDAC inhibition (Figure S2F). These results argue against the first proposed mechanism being responsible for the reduction in ATAC-seq signal at THS in TSA-treated samples. Rather, our findings suggest that genome-wide hyper-acetylation following HDAC inhibition encourages gains in accessibility at repeat elements.

We next sub-divided the repeat regions further by examining the changes in accessibility occurring at various classes of repetitive elements (Figure 2G). Interestingly, we found that the fraction of reads mapping to LINE elements increased upon TSA treatment although this was significant in DNMT.TKO but not WT cells. At LTR sites, both loss of DNA methylation and HDAC inhibition had an additive effect on accessibility. Differential analysis of the ATAC-seq coverage between samples revealed that there were significant increases in the number of ATAC-seq reads mapping to 33 of the 132 LINE-1 element subfamilies in TSA versus DMSO-treated wild-type cells (Figure S2G, Table S3). In DNMT.TKO cells, significant gains were detected for an additional five subfamilies (Figure S2H). Strikingly, it was the most conserved LINE-1 subfamilies that showed the most significant gains in accessibility in TSA-treated cells (Figure 2H), suggesting that the presence of intact transcription factor binding sites may be important in permitting accessibility. Amongst the LTR repeats, accessibility gains in DNMT.TKO cells predominantly occurred at ERVK-type elements (Figure S2I). HDAC inhibition potentiated these gains while also increasing accessibility at some ERVL and ERVL-MalR-type repeats.

Together, our ATAC-seq analysis indicates that a subset of normally methylated loci increase in accessibility in the absence of DNA methyltransferases, likely representing novel transcription factor binding sites. A distinct effect was observed upon HDAC inhibition, where the resulting genome-wide hyper-acetylation lead to an increase in accessibility outside of previously hyper-accessible sites (THS), particularly at a subset of LINE elements and LTR retrotransposons.

### DNA methylation and HDAC activity can modulate transcription factor occupancy

Elevated chromatin accessibility may be a result of abnormal TF binding (Domcke et al., 2015; Maurano et al., 2015), however it is not known whether our chromatin perturbations cause more widespread TF binding without inducing transposase hypersensitivity. Thus, we decided to directly measure how DNA methylation and histone acetylation affect the genomic distribution of selected transcription factors.

We performed ChIP-seq in WT and DNMT.TKO mESCs in the presence or absence of TSA, to map the genome-wide occupancy of five transcription factors (GABPA, MAX, NRF1, SP1 and YY1) that possess distinct DNA-binding domains and are known to bind cognate sequences containing CpG dinucleotides (Figure 3A).

**Figure 3:**
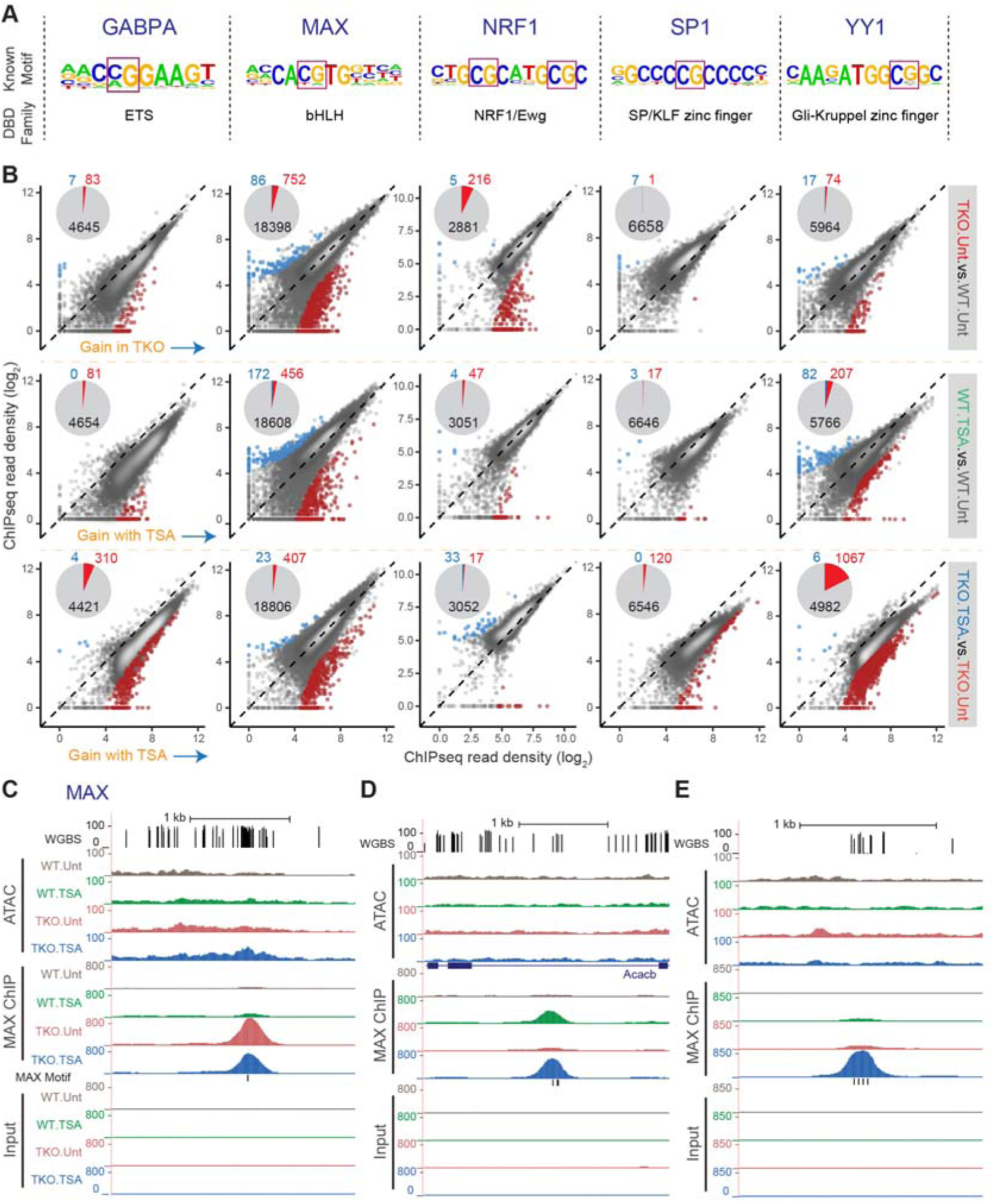
DNA methylation and HDAC activity can modulate transcription factor occupancy. **A.** For each transcription factor, the position weight matrix (PWM) of its known motif (as described in the HOMER database, Heinz et al., 2010) and its DNA-binding domain (DBD) type are shown. **B.** Pairwise comparisons of normalised GABPA, MAX, NRF1, SP1 or YY1 ChIP-seq signal for all identified occupancy peaks. Regions with significantly differential occupancy (fold change > 4 and adjusted p-value < 10^−3^) are coloured on the scatter plots and their numbers are summarised in the form of pie charts (light blue = significant decrease; red = significant increase). ChIP-seq signal from three biological replicate samples was averaged. **C.** Representative UCSC genome browser snapshot showing CpG methylation levels (Habibi et al., 2013), ATAC-seq, MAX ChIP-seq and Input ChIP-seq read coverage. The position of genes and that of sequences that match the MAX motif are shown. ATAC-seq and ChIP-seq coverage graphs were generated after merging alignments from replicate samples. Mm10 coordinates: chr6:35,364,240-35,368,757. **D.** Same as (C). Mm10 coordinates: chr5:114,231,923-114,236,836. **E.** Same as (C). Mm10 coordinates: chr7:44,781,271-44,782,839.

Using the DANPOS peak calling algorithm (Chen et al., 2013), 4735, 19236, 3102, 6666 and 6055 regions were identified as being reproducibly occupied by GABPA, MAX, NRF1, SP1 and YY1 in at least one experimental condition, respectively (Table S2). Biological replicates clustered together (Figure S3A) and the DNA motifs known to be bound by these transcription factors were the most highly enriched within the identified ChIP-seq peaks relative to the entire collection of motifs in the HOMER database, providing assurance of the specificity of these ChIP assays (Figure S3B).

For all five transcription factors, loss of DNA methylation or HDAC inhibition resulted in increased transcription factor occupancy (Figure 3B). However, the relative impact of these two alterations varied depending on the individual protein.

In accordance with our ATAC-seq analysis (Figure S2C), NRF1 gained novel binding sites in DNMT.TKO compared to wild-type cells (Figure 3B). As these regions are methylated in wild-type cells (Figure 4A), this suggests that DNA methylation restricts NRF1 binding at certain genomic loci, in line with previous reports (Domcke et al., 2015; Kumari and Usdin, 2001). On the other hand, HDAC inhibition did not have a substantial effect on the genome-wide occupancy of NRF1 with relatively few sites showing differential binding upon TSA treatment in wild-type or DNMT.TKO cells (Figures 3B and S3C). Moreover, few of the sites that appear to gain NRF1 occupancy after HDAC inhibition contain sequences that match its consensus recognition motif, suggesting that these binding events may be indirect or false positives (Figure 4B).

**Figure 4:**
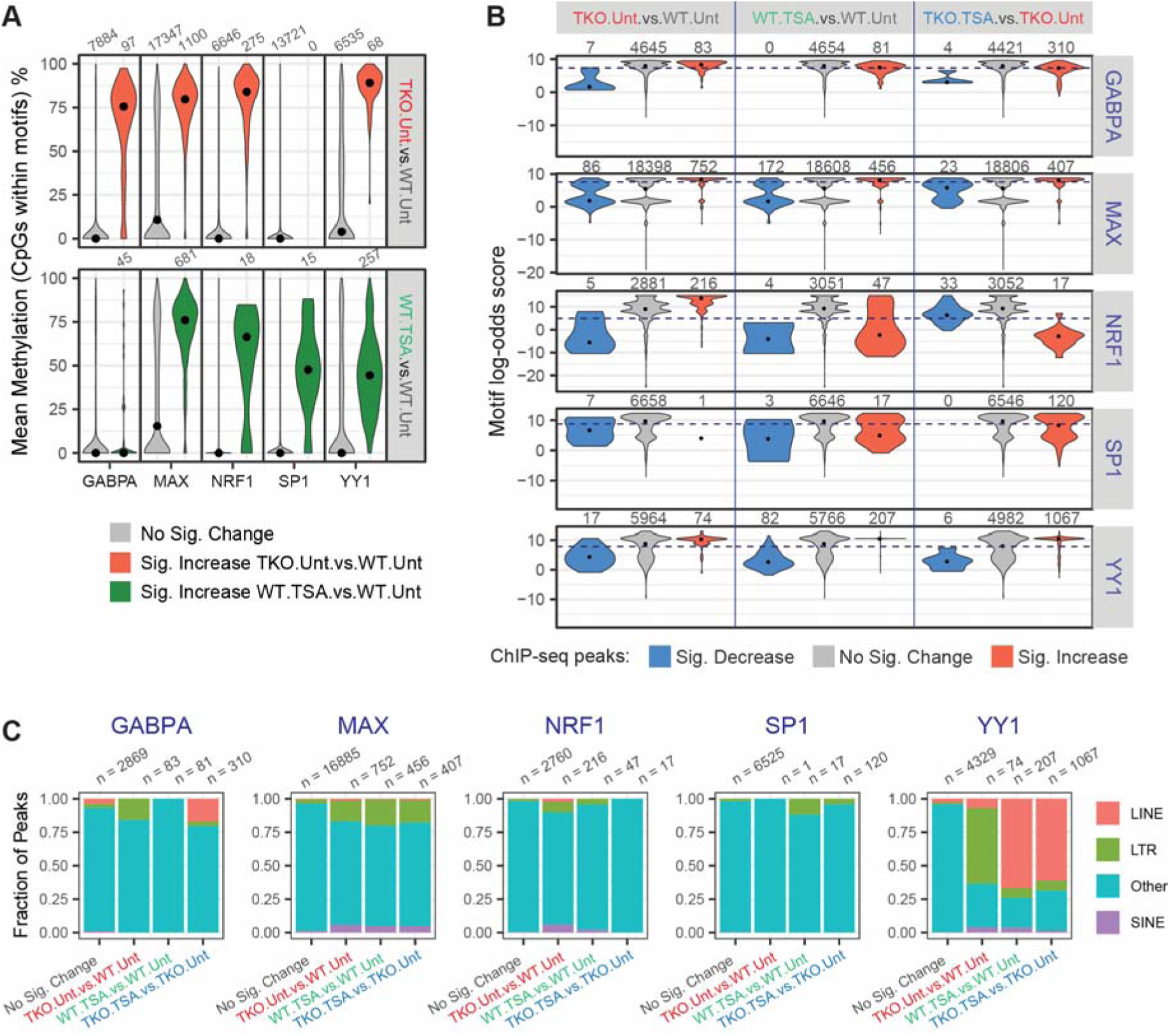
Characteristics of transcription factor binding sites. **A.** Distribution of CpG methylation levels within the transcription factor motifs that underlie ChIP-seq peaks. For each transcription factor, we isolated all sequences that were located within peak regions that matched the relevant position weight matrix and determined the methylation status of the CpG nucleotides. CpG sites are grouped according to their differential occupancy. Black dots indicate the median. **B.** Distribution of maximum log-odds scores found at different subsets of ChIP-seq peaks. Within each interval of the GABPA, MAX, NRF1, SP1 and YY1 peak-sets, the sequence most similar to the respective PWM was identified and its log-odds score, indicative of the deviation from the consensus sequence, was plotted. Black dots indicate the median. The dashed lines indicate the log-odds score threshold above which a sequence is said to match the PWM. **C.** Fraction of ChIP-seq peaks that overlap an annotated transposable element. ChIP-seq peaks are classified based on their differential occupancy.

Only few differential binding sites were identified for SP1 when comparing TSA-treated or DNMT.TKO cells to wild-type cells suggesting that this transcription factor’s occupancy is mostly unaltered by global loss of DNA methylation or genome-wide hyper-acetylation alone (Figure 3B). Nevertheless, we identify a 120 loci that gain SP1 binding upon HDAC inhibition in hypomethylated cells. Nearly half of those loci (48%) contained sequences highly similar to the GC box motif (Figure 4B).

Occupancy of GABPA, MAX and YY1 can be modulated by both DNA methylation and histone acetylation (Figure 3B). For these TFs, the occupancy peaks that appear after DNA methylation loss or HDAC inhibition predominantly harbour the respective TF binding motif (Figure 4B). Interestingly, for some of the novel binding sites, DNA methylation alone is sufficient to restrict occupancy of these factors (Figures S3C, 3C, S4A and S4D), whereas for others HDAC inhibition appears to be responsible for impeding binding (Figures S3C, 3D, S4B and S4E). Moreover, at a set of potential binding sites, both removal of DNA methylation and HDAC inhibition were necessary for transcription factor occupancy (Figures S3C, 3E, S4C and S4F).

The motifs present at DNMT.TKO-specific binding sites for GABPA, MAX and YY1 have elevated CpG methylation levels in wild-type cells (Figure 4A). It is therefore likely that DNA methylation is directly impeding the binding of these proteins to DNA, in agreement with previous *in vitro* measurements or locus-specific observations (Gaston and Fried, 1995; Kim et al., 2003; Lucas et al., 2009; Nickel et al., 1995; Satyamoorthy et al., 1993; Yin et al., 2017; Yokomori et al., 1995). Interestingly, the sites that gained GABPA occupancy upon HDAC inhibition, in wild-type cells were unmethylated (median CpG methylation of 0%), while those bound by MAX and YY1 under the same condition were more frequently methylated (76% and 44% median CpG methylation respectively, Figure 4A). The avoidance of methylated loci by GABPA suggests that this transcription factor might show the highest fraction of sites that significantly gain in occupancy when comparing DNMT.TKO cells treated with TSA to untreated wild-type cells. Indeed, GABPA occupancy significantly increased at 38% of all sites, while YY1 and MAX occupancy was gained at 9% and 21% of sites respectively (Figure S4G), demonstrating that both DNA methylation and histone acetylation influence the binding of these transcription factors to different degrees.

Although all five transcription factors predominantly occupy gene promoter regions (within 1500bp of a transcription start site (TSS)), the majority of ChIP-seq peaks that showed significant increases in occupancy upon DNA methylation loss or HDAC inhibition were situated at promoter distal genomic sites (Figure S5A). The exception to this is GABPA, for which over half of the sites displaying significant gains in occupancy in TSA-treated cells are located proximal to a TSS. This is likely related to our observation that TSA treatment relocates some of GABPA exclusively to non-methylated loci, which are often found within CpG islands near transcription start sites.

We further examined whether novel intergenic ChIP-seq peaks overlapped with transposable elements (LTRs, LINEs and SINEs). Most strikingly, over half of DNMT.TKO or TSA-specific YY1 ChIP-seq peaks overlapped with transposable elements (Figure 4C). Particularly, 652 out of 1067 YY1 binding sites acquired in the absence of DNA methylation and HDAC activity locate to LINE retrotransposons. More specifically, 537 of these overlapped with an L1Md_T type LINE repeat (82%) and 54 with an L1Md_Gf element (8.3%) (Figure S5B). Most L1Md_T and L1Md_Gf elements bound by YY1 in TKO.TSA samples contained at least one, sometimes an array of YY1 binding motifs within their 5’UTR regions (Figures S5C and S4E). Of note, LINE elements and particularly the L1Md_T and Gf subtypes also showed significant gains in accessibility in TSA treated cells (Figures 2G and 2H), consistent with a model in which transcription factor binding in the absence of HDAC activity and DNA methylation is contributing to chromatin opening.

A high fraction of the YY1 binding sites acquired in DNMT.TKO cells overlapped with LTR retrotransposons of which 34/42 were ERVK-type elements (Figure 4C). Interestingly, YY1 occupancy increases only modestly in the absence of DNA methylation or following HDAC inhibition (Figures S5D (left panel) and S4F). This differs from YY1 occupancy at LINE elements which occurs after HDAC inhibition but not as a consequence of DNA methylation loss alone (Figure S5D (right panel)). In both cases, YY1 occupancy is highest after HDAC inhibition in DNA methylation deficient cells (Figure S5D).

Altogether, our analysis of ChIP-seq data indicates that both DNA methylation and HDAC activity play distinct roles in limiting occupancy of each of the examined TFs to a fraction of their binding sites. Furthermore, we did not detect a significant loss in transcription factor occupancy at pre-existing peaks following HDAC inhibition, whereas novel binding events were observed throughout the genome. These findings support our previous interpretation of the ATAC-seq data: that chromatin accessibility increases preferentially outside of established THS and particularly at LINE and LTR repeat elements.

### Transcription factor occupancy can promote chromatin accessibility in mESCs with perturbed DNA methylation or HDAC activity

We next asked whether the gains in transcription factor occupancy in unmethylated or hyper-acetylated cells were directly associated with increased chromatin accessibility. For each transcription factor, we evaluated the changes in ATAC-seq signal at intervals that were occupied in one of our conditions. Generally, increased GABPA, MAX, NRF1 and YY1 binding in DNMT.TKO cells were associated with significant gains in accessibility (Figure 5A, top row). Novel binding events mostly occurred at sites that were previously inaccessible (Figure 5B, top) and a few coincided with an increase in ATAC-seq signal that was sufficient to be detected as a THS. Elevated transcription factor occupancy following HDAC treatment was also associated with significant gains in accessibility, although for GABPA this was only found to be the case after TSA treatment in DNMT.TKO but not WT cells (Figure 5A, bottom two rows). Whereas HDAC inhibition led to increased occupancy of MAX, NRF1 and YY1 at predominantly inaccessible loci (ie. that do not overlap any THS), the majority of sites to which GABPA and SP1 were relocated were previously accessible (Figure 5B, bottom). This finding is in agreement with our previous observations that all GAPBA and half of SP1 TSA-specific binding sites occur at promoter regions with low levels of DNA methylation (Figures 4A and S5A). Noticeably, novel TF binding events that appear following HDAC inhibition are less commonly associated with increased accessibility when compared to DNMT.TKO-specific binding events while the amplitude of change is also smaller (Figure 5A). Whilst this observation might reflect a different capacity of HDAC inhibition to induce THSs, compared to DNA demethylation, we cannot exclude the possibility that the lower signal is a consequence of the depletion in ATAC-seq reads from THSs after treatment with TSA.

**Figure 5:**
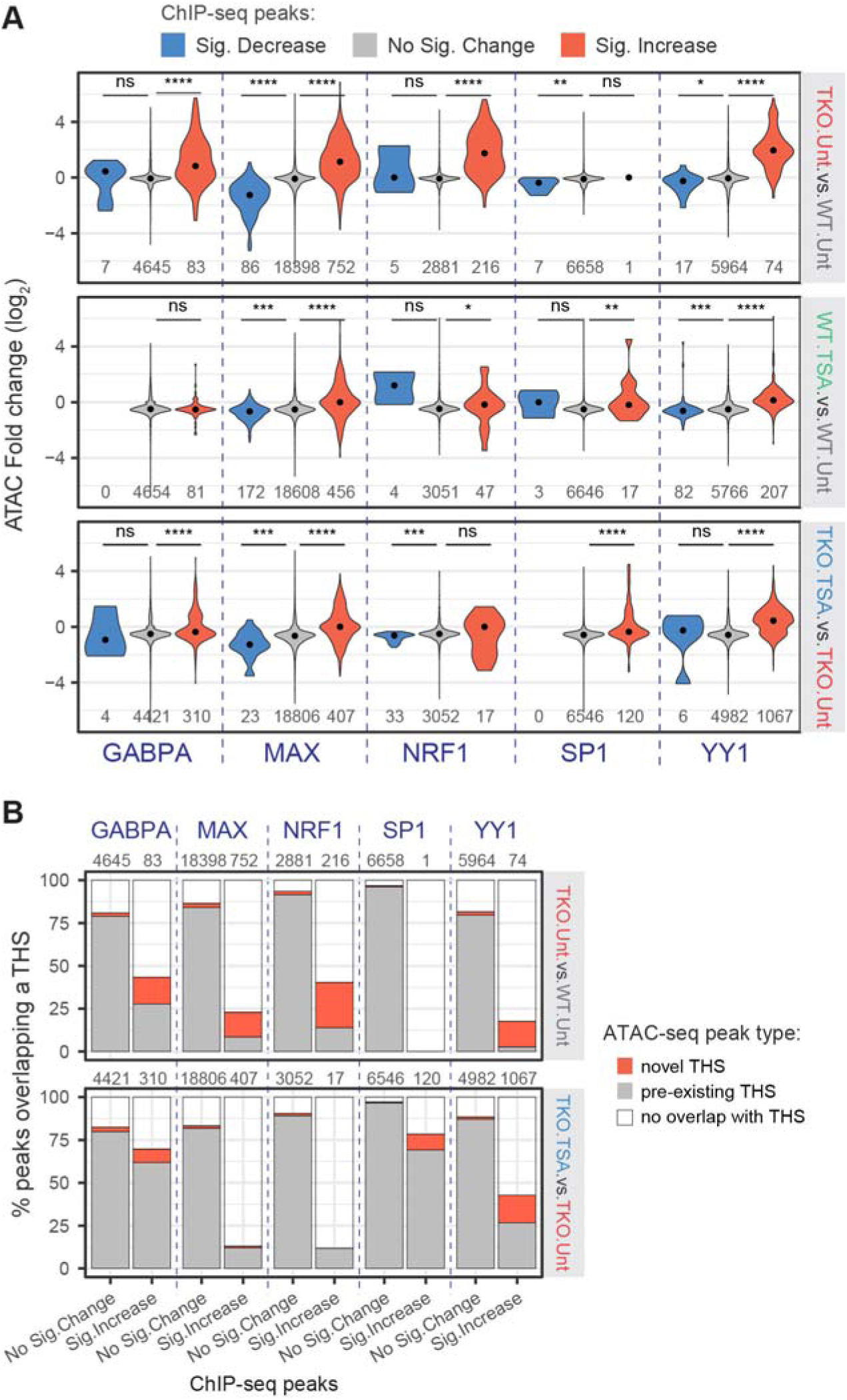
Transcription factor occupancy can promote chromatin accessibility in mESCs with perturbed DNA methylation or HDAC activity. **A.** Quantitation of ATAC-seq fold change at transcription factor ChIP-seq peaks grouped according to their differential occupancy. Black dots indicate the median ATAC-seq fold change. One-tailed Mann-Whitney U tests were used to determine significant differences in ATAC-seq fold change across peaksets: * p-value <0.05; ** p-value <0.01; *** p-value < 0.001; **** p-value < 10^−10^; *ns* = non-significant (p-value > 0.05). **B.** Percentage of ChIP-seq peaks that overlap with a THS region. In the top panel, a “pre-existing” THS refers to an ATAC-seq peak identified in samples generated from untreated WT cells while a novel THS is identified in DNMT.TKO but not WT cells. In the bottom panel, a “pre-existing” THS was identified in samples generated from untreated DNMT.TKO cells while a novel THS is identified in TSA-treated DNMT.TKO cells only.

### DNA methylation loss and HDAC inhibition affect the expression of a specific genes and repetitive elements

Having established that both DNA methylation loss and HDAC inactivation can lead to changes in chromatin accessibility and transcription factor occupancy, we next examined the consequences for gene expression. To this aim, we performed RNA-seq using wild-type and DNMT.TKO cells following treatment with TSA or DMSO. Loss of DNA methylation was associated with the differential expression of hundreds of genes (5.8% of total) whilst HDAC inhibition had a more profound effect, with 20% and 27% of genes being significantly deregulated in TSA-treated wild-type and DNMT.TKO cells respectively (Figure S6A, Table S4).

We first explored the relationship between changes in chromatin accessibility and gene expression. To this aim, we assigned each ATAC-seq peak to its nearest transcription start site and examined the gene expression differences. We observed weak correlations between changes in chromatin accessibility and expression of the neighbouring gene (Spearman Correlation Coefficients for TKO.DMSO versus WT.DMSO, WT.TSA versus WT.DMSO or TKO.TSA versus TKO.DMSO are 0.15, 0.11 and 0.10 respectively) (Figure 6A), indicating that most chromatin accessibility alterations are not associated with transcriptional changes at the closest gene. Nevertheless, more differentially accessible THSs show matching changes in neighbouring gene expression than expected by chance (Figure 6A). Many of the differential THSs are promoter distal and may represent loci with no regulatory function or that regulate transcriptional elements other than the closest gene. Limiting the list of ATAC-seq peaks to those that are promoter proximal (within 1500bp of a TSS) strengthens the correlation with gene expression when comparing WT and DNMT.TKO cells, but not TSA and control treatments (Figure S6B). At 28 promoters, significant changes in accessibility did associate with significantly increased gene expression in DNMT.TKO cells (Figures 6B, 6C and S6B). Notably, these 28 genes are enriched for genes whose expression is normally restricted to testes (DAVID functional annotation analysis, Table S5) fitting with a known role for DNA methylation in controlling the expression of germ-cell specific genes (Auclair et al., 2014; Borgel et al., 2010; De Smet et al., 1999; Fouse et al., 2008; Maatouk et al., 2006). In conclusion, DNA methylation can restrict gene expression by maintaining an inaccessible chromatin state; however the majority of gained THSs both in de-methylated or hyper-acetylated genomes are unable to affect the mRNA abundance of the nearest genes.

**Figure 6:**
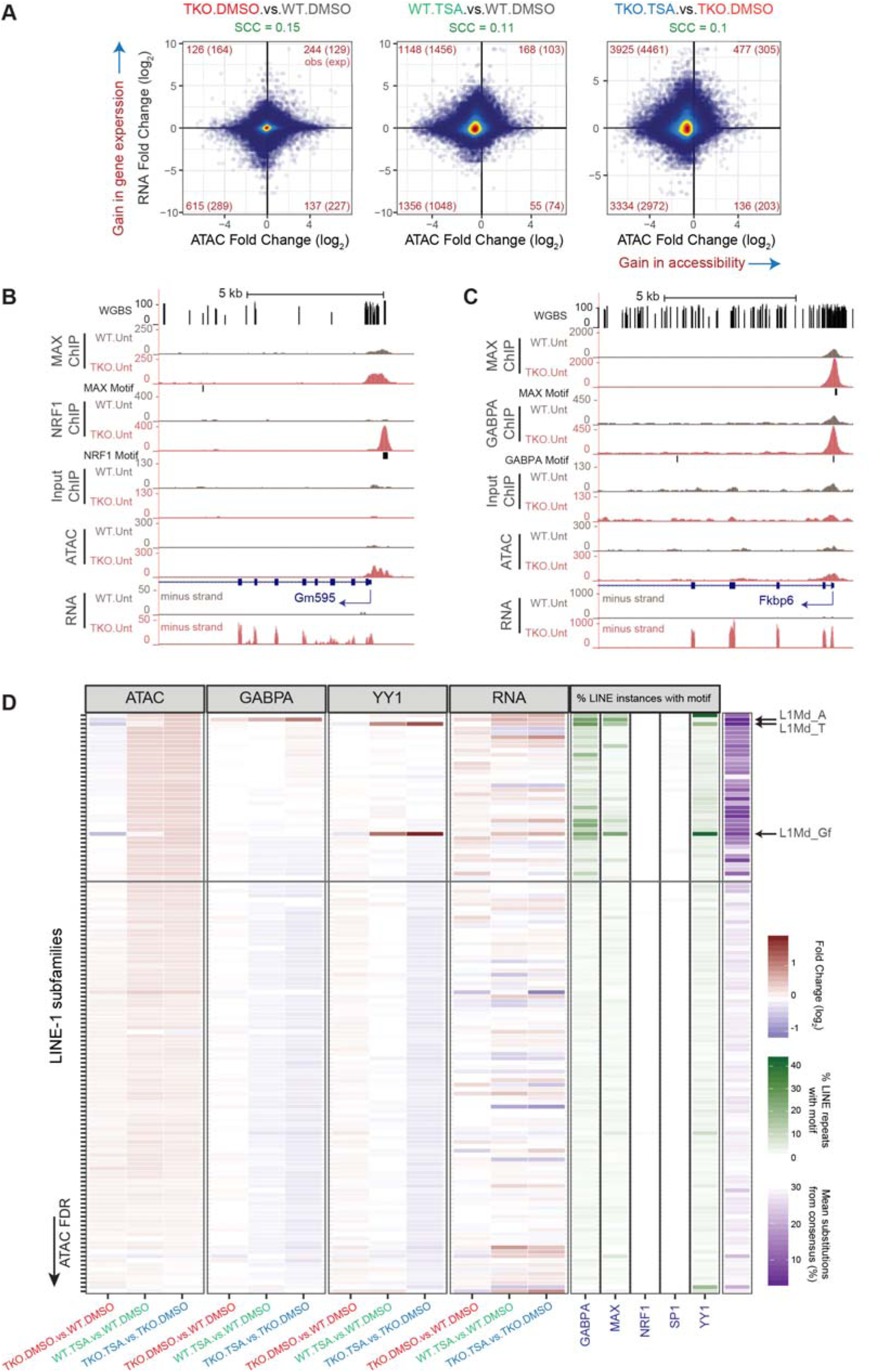
DNA methylation loss and HDAC inhibition affect the expression of a specific genes and repetitive elements. **A.** Changes in accessibility at every THS region were compared to changes in expression at their closest gene. The analysis was performed on 60052 THS:gene pairs involving 15298 genes. The numbers of ATAC-seq peaks associated with significant changes in both accessibility <u>and</u> gene expression are indicated in red in each quadrant. In brackets are indicated the expected number of sites showing both significant changes in accessibility and gene expression based on the total number of significant differential events. SCC = Spearman Correlation Coefficient. **B.** Representative UCSC genome browser snapshot showing CpG methylation levels (Habibi et al., 2013), ChIP-seq, ATAC-seq and strand-specific RNA-seq read coverage. The position of genes and that of sequences that match the TF motifs are shown. ATAC-seq, ChIP-seq and RNA-seq coverage graphs were generated after merging read counts from replicate samples. Mm10 coordinates: chrX:48,867,013-48,884,315. **C.** Same as (B). Mm10 coordinates: chr5:135,334,899-135,352,814. **D.** For each LINE-1 subtype (N = 132), we plotted the fold change in ATAC-seq, ChIP-seq or RNA-seq signal along with scores relating to their sequence conservation (purple) or the presence of selected TF binding motifs (green). LINE-1 subtypes were sorted based on ATAC-seq false discovery rate (FDR) when comparing TSA- to DMSO-treated DNMT.TKO cells. See also Table S3.

Considering our observations that certain subtypes of LINE-1 and LTR retrotransposons were selectively accessible and bound by transcription factors following DNA methylation loss or HDAC inhibition, we turned our attention to transcription originating from these repetitive elements. In the case of LINE-1 and ERVL-type LTR repeats, we noticed that increases in accessibility that are observed following HDAC inhibition but not after loss of DNA methylation were associated with elevated levels of RNA originating from various subtypes (Figure 6D and S6C). These included the highly conserved L1Md_T, Gf and A subtypes for which there was also enrichment for YY1 or GABPA ChIP-seq reads (Figure 6D), as well as MT2-Mm and MERVL-int ERVL elements. In the case of ERVK-type repeats, loss of DNA methylation rather than HDAC inhibition accounts for the most pronounced increases in accessibility although this is restricted to a subset of repeat types, including IAP elements (Figure S6D). However, the expression from ERVK-type elements is most dramatically induced when both DNA methylation and HDAC activity are abolished (comparing TKO.TSA cells to WT.DMSO). This observation agrees with previous reports documenting a synergistic activation of ERVK-type repeats in response to a combined removal of DNA methylation and HDAC activity (Brocks et al., 2017; Cameron et al., 1999; Coffee et al., 1999; Walter et al., 2016).

In summary, elevated transcription factor binding and genome accessibility in the absence of DNA methylation or after HDAC inhibition promote activation of small subset of genes, whilst up-regulating expression of LINE-1 and LTR repeats.

## Discussion

In this study, we examined the roles of DNA methylation and histone deacetylase activity in genome accessibility, transcription factor occupancy and gene regulation. Our data indicates that transcription factors have different abilities to sense chromatin modifications. Even when examining a small number of TFs, we were able to capture distinct scenarios in which TF occupancy is affected by neither DNA modifications nor histone acetylation (SP1), exclusively by DNA methylation (NRF1), or by both (GABPA, MAX, SP1 and YY1) (Figure 7).

**Figure 7:**
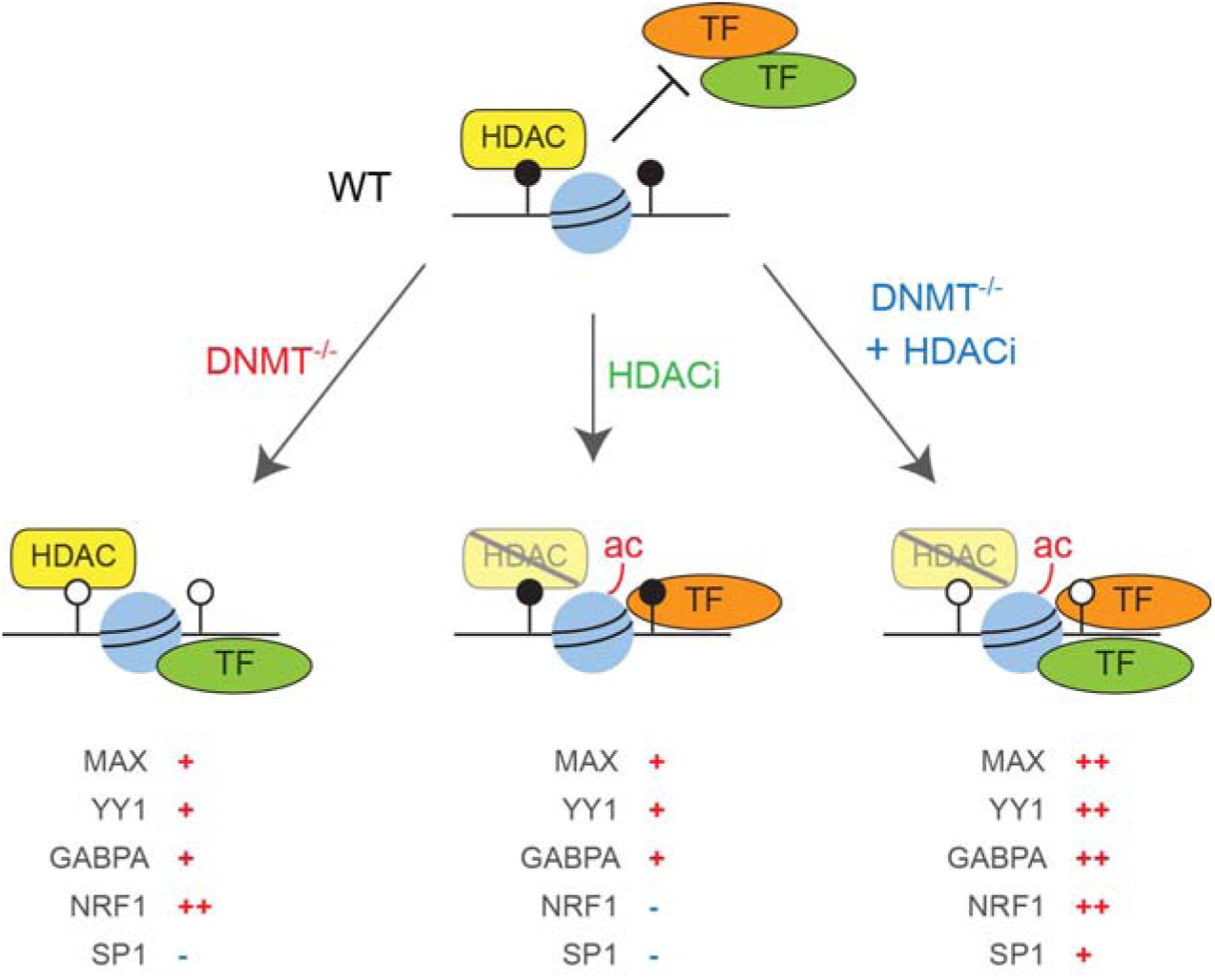
Summary model illustrating impact of DNA methylation and HDAC inhibition on transcription factor occupancy.

In contrast to previous studies, which discovered altered TF occupancy by identifying enriched TF binding motifs from chromatin accessibility data, we directly measured occupancy of selected candidate TFs. The accuracy of using genome accessibility data to infer TF displacement depends on, firstly, the ability of aberrant binding events to induce a change in accessibility and, secondly, gaining enough accessible sites per TF to reach statistical power for the identification of binding motifs. Using direct measurements of occupancy, we demonstrate instances where TF mis-localisation does not cause different genome accessibility. For example, in DNA methylation deficient cells ∼75% of novel MAX binding peaks do not overlap with THSs. Thus TF occupancy is likely to be more sensitive to chromatin structure than previously thought.

The total number of new binding events varies between transcription factors and does not correlate with the number of all potential binding sites (i.e. sequences that match its cognate motif perfectly). For example, in DNA methylation-deficient cells NRF1 gains occupancy at 216 loci (increase by 0.6% of all potential binding sites), whereas MAX gains occupancy at 752 (0.26%), YY1 – 74 (0.02%) and GABPA – 83 (0.02%) loci. Changes of a similar magnitude have been reported independently for another two proteins MYC-N and CTCF in DNMT.TKO mES cells (Stadler et al., 2011; Yin et al., 2017). As the NRF1 motif is the most prevalent motif underlying DNMT.TKO-specific chromatin hyper-accessible sites based on our ATAC-seq data and DNase-seq data produced in an analogous DNMT.TKO cell line (Domcke et al., 2015), it is unlikely that any other transcription factor undergoes a more extensive genomic redistribution following DNA methylation depletion in mouse ES cells. Comprehensive *in vitro* experiments indicate that there are over a hundred TFs whose affinity to DNA is diminished by cytosine methylation (Hu et al., 2013; Yin et al., 2017). Therefore, while the contribution of DNA methylation in restricting the occupancy of individual transcription factors appears limited, the cumulative effect of DNA methylation loss on the gene regulatory network is likely to be more substantial. Moreover, the contribution of DNA methylation to TF occupancy may diverge or be more important in cell types other than mouse embryonic stem cells, which are unique in their capacity to maintain homeostasis in its absence. For example, genomic CTCF occupancy increases by 5.4% in hypomethylated compared to wild-type HCT116 cells (Maurano et al., 2015) but shows almost no difference in DNMT.TKO versus wild-type mouse ES cells other than at several imprinted loci (Stadler et al., 2011). Thus aberrant TF binding in the absence of DNA methylation is likely to contribute to the essential nature of DNMT enzymes in differentiation and during development (Li et al., 1992; Liao et al., 2015; Okano et al., 1999; Tsumura et al., 2006; Tucker et al., 1996).

Unlike the effect of DNA methylation depletion, HDAC inhibition did not result in the emergence of many discrete Tn5 hypersensitive loci. Rather, increased chromatin accessibility was observed throughout the genome and particularly at a subset of abundant LINE and LTR transposable elements that led to a significant skew in the composition of ATAC-seq libraries. Despite genome-wide hyper-acetylation, aberrant binding of the examined transcription factors occurred at a scale similar to that seen following the loss of DNA methylation. GABPA, MAX, NRF1 and YY1, but not SP1, were sensitive to DNA methylation, whereas only MAX, GABPA and YY1, but not NRF1, displayed increased genomic occupancy upon HDAC inhibition in wild-type cells. Notably, perturbations to the DNA methylation machinery or HDAC activity had distinct impacts on MAX, GABPA and YY1 localisation both in terms of the number of aberrantly occupied loci as well as the genomic location of these alterations, indicating that DNA methylation does not depend on HDAC activity to restrict TF occupancy in mouse ES cells. One of the models that is recurrently proposed to explain the transcriptional silencing activity of DNA methylation suggests that methylated CpG sequences recruit MBD-domain proteins whose associated HDAC enzymes promote transcriptional repression (Eden et al., 1998; Kokura et al., 2001; Lyst et al., 2013; Nan et al., 1998). However, we show that the effect of HDAC inhibition on chromatin accessibility, TF occupancy and gene expression does not recapitulate that of DNA methylation loss, in line with similar observations in mESCs (Brunmeir et al., 2010; Reichmann et al., 2012) and other cell lines (Brocks et al., 2017; Lorincz et al., 2000) arguing that, at large, DNA methylation functions independently of HDACs. A more direct role for cytosine methylation in lowering the affinity between transcription factors and their cognate DNA sequences is however consistent with our data, as well as being supported by various biochemical and structural experiments (Dantas Machado et al., 2015; Prendergast and Ziff, 1991; Spruijt et al., 2013; Stephens and Poon, 2016; Yin et al., 2017).

On the other hand, the manner by which histone deacetylation hinders transcription factor binding is unclear and remains to be elucidated. *In vitro* data indicate that nucleosomes made up of histones with acetylated N-terminal tails associate more loosely with DNA and are less prone to form compact higher order chromatin structures (Anderson et al., 2001; Hong et al., 1993; Lee et al., 1993; Wang and Hayes, 2008). Alone, this could explain the increased ATAC-seq signal that accompanies histone hyper-acetylation outside of THS sites in mESC cells that we have treated with TSA (Vidali et al., 1978). Moreover, such a weakening of the interaction between histones and DNA may be permissive to the binding of TFs that usually have to compete with nucleosomes for binding to DNA (Hayes and Wolffe, 1992; Polach and Widom, 1995; Raveh-Sadka et al., 2012). Indeed, most TFs preferentially bind to nucleosomal-free DNA, as is the case for multiple members of the bHLH, ETS and Znf_C2H2 DNA binding domain families to which MAX, GABPA and YY1 belong, respectively (Zhu et al., 2018). One can also envision that hyper-acetylation permits TF binding through indirect mechanisms, such as ones involving intermediate chromatin remodelling factors that possess BRD domains (Fujisawa and Filippakopoulos, 2017), or by obstructing the methylation of lysine residues such as H3K9 or H3K27 that are known to promote the formation of chromatin conformational states that are inhibitive to TF binding through the recruitment of HP1, KAP1 or polycomb-group proteins (Bannister et al., 2001; Canzio et al., 2011; Francis et al., 2004; Lachner et al., 2001; Larson et al., 2017; Matoba et al., 2014; Soufi et al., 2012; Strom et al., 2017). In support of the latter suggestion, certain repeat elements such as MERVL, LINE and IAP retrotransposons that we show are upregulated upon HDAC inhibition are also activated following the disruption either of the H3K9 methyltransferases SETDB1, SUV39H and G9a/GLP or the KAP1 repressor protein (Karimi et al., 2011; Macfarlan et al., 2012; Maksakova et al., 2013; Reichmann et al., 2012).

While HDAC activity and DNA methylation may act as individual barriers to different transcription factors, their activity does collaborate to regulate specific genomic elements. Indeed, at certain loci, including at a subset of ERVK LTR elements (including IAP elements), transcription factor occupancy was only detected upon both DNA methylation loss and HDAC inhibition. Several single copy genes and transposable elements underwent pronounced transcriptional activation in this condition, while they were mostly unaffected by either treatment alone. Such synergistic activity of DNA methylation loss and HDAC inhibition has previously been described at LTR12-type transposable elements in human lung cancer cells (Brocks et al., 2017) and at several methylated CpG island genes in a colorectal carcinoma cell line (Cameron et al., 1999). Together, these data indicate that DNA methylation and HDAC activity act independently to impede TF occupancy, such that the presence of one is sufficient to compensate for the absence of the other. These synergistic effects could be explained in several ways. The first possibility is that an individual protein is sensitive to DNA methylation and histone acetylation, only being able to bind (and promote transcription) in the absence of both. An alternative model is that stable TF-DNA complexes and transcriptional activity can only occur through the cooperative binding of multiple factors, among which some have intrinsic sensitivity to DNA methylation (e.g. NRF1) and others to histone acetylation (e.g. YY1) (Figure 7). For example, different sets of sites are occupied by MAX following TSA treatment or DNA methylation loss, which might reflect sets of loci at which the occupancy of MAX is facilitated by distinct co-factors that are dependent on histone acetylation or DNA hypomethylation, respectively. Indeed, MAX is known to form heterodimers with diverse protein partners (Blackwood et al., 1992; Hurlin et al., 1995, 1997, 1999) and it is conceivable that these different complexes have different affinities for chromatin modifications. C-Myc, for instance, has a strong preference for binding E-box motifs that are in regions marked with H3K4 methylation and H3K27 acetylation (Guccione et al., 2006; Sabò and Amati, 2014).

Although there are only a few genomic loci for which HDAC inhibition or DNA methylation loss appears to be directly responsible for altered TF binding and transcriptional activity, these elements are of biological and potentially clinical significance. LINEs and ERVLs for example, are reported to be essential for early embryonic development following fertilisation (Jachowicz et al., 2017; Kruse et al., 2019; Percharde et al., 2018), a period during which histone acetylation is elevated and DNA methylation is low (Eckersley-Maslin et al., 2016; Ishiuchi et al., 2015; Macfarlan et al., 2012). In somatic tissues, aberrant expression of germ-cell specific genes and retrotransposons is likely to be detrimental. Inhibitors of HDAC and DNMT enzymes are being used for the treatment of acute myelogenous leukaemia and their use against solid tumours is being considered in combination with immune checkpoint inhibitors (Daskalakis et al., 2018; Navada et al., 2014; West and Johnstone, 2014). Several reports suggest that aberrant RNA transcripts or their translated peptides produced following DNA hypo-methylation and/or HDAC inhibition may be recognised by cytosolic or extracellular components of the innate immune system, thus contributing to the efficacy of these drugs (Brocks et al., 2017; Chiappinelli et al., 2015; Haffner et al., 2018; Roulois et al., 2015; Stone et al., 2017).

In summary, we demonstrate that deacetylation of histones and DNA methylation are two important activities in chromatin, which collaborate to limit TF occupancy and contribute to silencing of repeat expression.

## Acknowledgments

S.K. is supported by the Ludwig Institute for Cancer Research. S.K. also acknowledges funding BBSRC BB/M001873/1 and Conrad N. Hilton Foundation. M.C. was funded by the Medical Research Council. We thank Masaki Okano for *Dnmt* TKO ESCs. We thank Hiromi Tagoh, Thomas Milne and Mary Muers for critical reading of the manuscript.

## Author Contributions

M.C. and S.K. conceived the study and wrote the manuscript with contribution from all authors; M.C. performed all experiments and data analysis, except as indicated below; H.W.K. and M.C. under supervision by R.J.K. and S.K performed ATAC-seq experiments; P.S. under supervision by S.K. and B.M.K. carried out LC-MS measurements.

## Declaration of Interests

The authors declare no competing interests.

## Materials and Methods

### Cell culture and treatment

*Dnmt1*^-/-^, *Dnmt3a*^-/-^ and *Dnmt3b*^-/-^ TKO (DNMT.TKO, (Tsumura et al., 2006)) and wild-type J1 male mouse embryonic stem cells were grown on 0.1% gelatin-coated dishes in DMEM (Lonza) complemented with 10% (v/v) Fetal Bovine Serum (Biosera, FB-1001G/500, lot #11484), 2 mM L-glutamine (Life Technologies), 1% (v/v) non-essential amino acids (Life Technologies), 1% (v/v) penicillin-streptomycin (Life Technologies), 10 ng/ml Leukaemia Inhibitory Factor (LIF, produced in-house following a protocol by Tomala et al., 2010) and 0.1 mM beta-mercaptoethanol (Sigma). Chicken DF-1 (Doug Foster strain 1) fibroblast cells were grown in DMEM (Lonza) supplemented with 10% (v/v) Fetal Bovine Serum (Biosera, FB-1001/500 lot #014BS386), 1% (v/v) penicillin-streptomycin (Life Technologies), 2 mM L-glutamine (Life Technologies) and 0.15% sodium bicarbonate (Life Technologies). All cells were incubated in a humidified incubator set at 37 °C and 5% atmospheric CO_2_.

HDAC inhibition in mESCs was achieved using Trichostatin A (TSA, Sigma, T1952). Cells were seeded 4-6 h prior to treatment; once attached their growth medium was replaced with medium containing 5 ng/µl TSA. TSA-containing media was refreshed after 24 h and cells were harvested for downstream applications after 36 h. Control DMSO-treated cells were seeded and treated in a similar manner however medium contained 0.0033% DMSO.

### Quantification of global 5mC by Mass Spectroscopy

Genomic DNA was extracted from mouse ES cells using the GeneJET Genomic DNA Purification Kit (ThermoFisher Scientific) following the manufacturer’s instructions. Hydrolysis was carried out in 150 µl reactions that contained 500 ng DNA, 100 mM NaCl, 20 mM MgCl2, 20 mM Tris pH 7.9, 1000 U/ml Benzonase, 600 mU/ml Phosphodiesterase I, 80 U/ml Alkaline phosphatase, 36 µg/ml EHNA hydrochloride and 2.7 mM deferoxamine for 2 hours at 37 °C. Following lyophilisation, nucleosides were resuspended in buffer A (10 mM ammonium acetate, pH 6) with 12.5 nM standards of +3 Da mdC and hmdC.

For the analysis by HPLC– QQQ mass spectrometry, a 1290 Infinity UHPLC was fitted with a Zorbax Eclipse plus C18 column, (1.8 µm, 2.1 mm 150 mm; Agilent) and coupled to a 6495a Triple Quadrupole mass spectrometer (Agilent Technologies) equipped with a Jetstream ESI-AJS source. The data were acquired in dMRM mode using positive electrospray ionisation (ESI1). mdC and hmdC were quantified by mass spectrometry, whereas dC was quantified by HPLC-UV. The gradient used to elute the nucleosides started by a 5 min isocratic gradient composed of 100% buffer A and 0% buffer B (100% methanol) with a flow rate of 0.4 ml/min and was followed by the subsequent steps: 5-8 min, 94.4% A; 8–9 min, 94.4% A; 9–16 min 86.3% A; 16–17 min 0% A; 17– 21 min 0% A; 21–24.3 min 100% A; 24.3–25 min 100% A. The AJS ESI settings were as follows: drying gas temperature 230 °C, the drying gas flow 14 lmin^−1^, nebulizer 20 psi, sheath gas temperature 400 °C, sheath gas flow 11 lmin^−1^, Vcap 2,000 V and nozzle voltage 0 V. The iFunnel parameters were as follows: high pressure RF 110 V, low pressure RF 80 V. The fragmentor of the QQQ mass spectrometer was set to 380 V and the delta EMV set to +200.

The raw mass spectrometry data was analysed using the MassHunter Quant Software package (Agilent Technologies, version B.08.01). The transitions and retention times used for the characterization of nucleosides and their adducts are summarized in Table S6. For the identification of compounds, raw mass spectrometry data was processed using the dMRM extraction function in the MassHunter software.

### Isolation of histone proteins and immunoblotting

Unless stated otherwise, all steps were performed on ice, spins performed in a centrifuge chilled to 4 °C and rotations set up in a cold room. Cells were lysed in lysis buffer (PBS supplemented with 0.5% (v/v) Triton-X100, 1x cOmplete^TM^ EDTA-free Protease Inhibitor Cocktail (Roche) and 10 mM sodium butyrate) for 10 min after which nuclei were washed once with lysis buffer. Histone proteins were solubilised by incubating the nuclei in 0.2 M H_2_SO_4_ with rotation overnight and collected by spinning at 16,000 × g for 10 min. The histone proteins were subsequently precipitated by adding trichloroacetic acid (TCA) to a final concentration of 25% and incubating them for at least 2 hours on ice followed by the centrifugation at 16,000 × g for 10 min. Following two washes with ice-cold acetone, histone proteins were resuspended in 1x SDS-PAGE Loading Buffer (40 mM Tris-HCl pH 6.8, 80 mM DTT, 1.6% SDS, 0.08% bromophenol blue, 8% glycerol) and denatured at 95 °C for 5 min. Samples were resolved on 15% SDS-PAGE gels and analysed by immunoblotting. Histone proteins were detected by using a rabbit polyclonal anti-pan-acetyl H3/H4 antibody (Upstate, 06-866; 1:5000 dilution), a rabbit polyclonal antibody anti-pan-acetyl H3 antibody (Millipore, 06-599; 1:2000 dilution) and a rabbit polyclonal antibody against total unmodified H3 (Abcam, ab1791; 1:40000 dilution). The Santa Cruz sc-2004 goat anti-rabbit HRP-conjugated antibody was used as a secondary antibody (1:10000 dilution).

### Calibrated Native Chromatin Immunoprecipitation with exogenous spike-in

Chicken DF-1 and mouse ES cells were harvested by trypsinisation. For each experimental condition, 5 × 10^6^ chicken DF-1 cells were added to 5 × 10^7^ mESCs. Cell mixtures were pelleted and resuspended in 1 ml ice-cold RSB Buffer (10 mM NaCl, 10 mM Tris (pH 8.0), 3 mM MgCl_2_). Chromatin was fragmented by adding 200 units of Micrococcal Nuclease (MNase, Fermentas) and incubating at 37 °C for 5 min. The reaction was stopped by the addition of 8 µl of 0.5M EDTA and spun at 2348 × g for 5 min at 4°C. The supernatant containing soluble nucleosomes was collected (S1 fraction). Nucleosomes in the remaining insoluble fraction were extracted by rotating in 300 µl Nucleosome Release Buffer (10 mM Tris pH 7.5, 10 mM NaCl, 0.2 mM EDTA, with 1x cOmplete^TM^ EDTA-free Protease Inhibitor Cocktail (Roche) and 10 mM sodium butyrate) for 1 h at 4 °C followed by the passing through a 25G needle five times. The supernatant was collected as described above and pooled with the S1 chromatin fraction. DNA fragment size was checked by agarose gel electrophoresis indicating that the majority of the fragments are mononucleosomal DNA (150-200 bp) with a small fraction of di- and tri-nucleosomal.

Immunoprecipitations were set up with 100 µl solubilised chromatin (roughly equivalent to 4 × 10^6^ cells) and 3 µg of an anti-acetylated histone H3 antibody (Millipore, 06-599) in a total of 1 ml Native ChIP Incubation Buffer (10 mM Tris pH 7.5, 70 mM NaCl, 2 mM MgCl_2_, 2 mM EDTA and 0.1% Triton X-100, 1x cOmplete^TM^ EDTA-free Protease Inhibitor Cocktail and 10 mM sodium butyrate) and rotated overnight at 4°C. Immune complexes were captured using 20 µl Protein A agarose beads (Roche, previously saturated with 1 mg/ml BSA and 1mg/ml yeast tRNA) for 1 hour at 4°C with rotation, washed four times with Native ChIP Wash Buffer (20 mM Tris pH 7.5, 2 mM EDTA, 125 mM NaCl, 0.1% Triton-X100) and once in TE buffer (10 mM Tris HCl pH 8.0, 1 mM EDTA) and subsequently eluted in 100 µl elution buffer (1% SDS, 0.1 M NaHCO_3_) by vigorous shaking for 30 min at room temperature. DNA was purified using the ZymoResearch ChIP DNA clean and concentrator kit (Zymo Research) following the manufacturer’s instructions. For input samples, DNA was purified from 100 µl of solubilised chromatin using the same DNA purification procedure.

Material was quantified using Qubit (Invitrogen) and the size profile analysed on the 2200 or 4200 TapeStation (Agilent, dsDNA HS Assay). Automated library preparation was performed on 5 ng input material using the Apollo prep system (Wafergen, PrepX ILMN 32i, 96 sample kit) and standard Illumina multiplexing adapters following manufacturer’s protocol up to pre-PCR amplification. Libraries were PCR amplified (18 cycles) on a Tetrad (Bio-Rad) using the NEBNext High-Fidelity 2X PCR Master Mix (NEB) and in-house single indexing primers (Lamble et al., 2013). Equivalent amounts of individual libraries were pooled. Paired-end sequencing was performed using a HiSeq4000 75 bp platform (Illumina, HiSeq 3000/4000 PE Cluster Kit and 150 cycle SBS Kit), generating a raw read count of >30 million reads per sample.

### ATAC-seq sample preparation with exogenous spike-ins

Assay for Transposase Accessible Chromatin (ATAC)-seq experiments were carried out by adapting the protocol published by (Buenrostro et al., 2013). To allow for calibration of the ATAC-seq signal between different samples, 5 × 10^5^ chicken DF-1 cells were spiked-in with 5 × 10^6^ mouse ES cells from each experimental condition. Cell mixtures were lysed in 1 ml of Lysis Buffer (50 mM KCl, 10 mM MgSO_4_x7H_2_O, 5 mM HEPES, 0.05% NP40 (IGEPAL CA630), 1 mM PMSF and 3 mM DTT) by incubating at room-temperature for 1 min followed by the two washes in 1 ml ice-cold RSB Buffer (10 mM NaCl, 10 mM Tris (pH 7.4), 3 mM MgCl_2_).

ATAC reactions were performed in 50 µl reactions in the presence of Tn5 reaction buffer (10 mM TAPS, 5 mM MgCl_2_ and 10% dimethylformamide), 5 × 10^4^ nuclei (resuspended in H_2_O) and 2 µl of 12 μM Tn5 transposase (produced in house following a protocol by Picelli et al., 2014) for 30 min at 37°C. DNA was purified using the QiaQuick MinElute columns (Qiagen) following manufacturer’s instructions.

Input samples were prepared by subjecting genomic DNA, which had been isolated from each sample prior to the ATAC reaction, to Tn5 tagmentation in reactions containing 5 ng genomic DNA, 1 µl of Tn5 and incubated at 55 °C for 7 min.

ATAC-seq libraries were prepared by 6-12 cycles of PCR amplification using custom made Illumina barcodes in the NEBNext Ultra II Q5 2x PCR Master Mix. PCR amplicons were purified and size selected using home-made SPRI beads (Rohland and Reich, 2012) in two rounds (1:0.6 then 1:1.4 DNA:beads ratio). ATAC-seq libraries were quantified using the NEBNext Library Quant Kit and pooled prior to the sequencing on an Illumina HiSeq4000 instrument at the Oxford Genomics Centre to generate 75 bp paired-end reads. ATAC-seq experiments were performed on four biological replicates.

### Chromatin immunoprecipitation for transcription factors

Chromatin was prepared from 1 × 10^8^ mouse ES cells grown to 70-90% confluency. Cells were crosslinked at room temperature for either 10 min in 1% methanol-free formaldehyde (Thermo Scientific) or 2 mM disuccinimidyl glutarate (DSG) for 1 hour followed by 10 min with 1% formaldehyde (YY1 only). Crosslinking reactions were quenched by the addition of glycine to a concentration of 200 mM. Chromatin was prepared at 4 °C by rotating the cells in 10 ml buffer LB1 (50 mM HEPES KOH pH 7.9, 140 mM NaCl, 1 mM EDTA, 10% Glycerol, 0.5% NP40, 0.25% Triton x-100, with 1x cOmplete^TM^ EDTA-free Protease Inhibitor Cocktail (Roche)), 10 ml buffer LB2 (10 mM Tris-HCl pH 8.0, 200 mM NaCl, 1 mM EDTA, 0.5 mM EGTA, with 1x cOmplete^TM^ EDTA-free Protease Inhibitor Cocktail) and 1 ml buffer LB3 (10 mM Tris-HCl pH 8.0, 100 mM NaCl, 1 mM EDTA, 0.5 mM EGTA, 0.1% sodium deoxycholate, 0.5% N-lauroylsarcosine, 1x cOmplete^TM^ EDTA-free Protease Inhibitor Cocktail). Chromatin was sheared in a 1 ml milliTUBE with AFA Fiber (Covaris) using a Covaris S220 AFA ultrasonicator (set at 150 W Peak Incident Power, 8% Duty Factor, 200 Cycles/Burst) for 30 min. Following sonication, the sheared chromatin was spun at 15,000 × g for 20 min and the soluble fraction collected and aliquoted for immediate use or storage at −80 °C.

Immunoprecipitation (IP) was performed using the sheared chromatin from approximately 1 × 10^7^ cells in Chromatin Dilution Buffer (16.7 mM Tris pH 8.1, 167 mM NaCl, 0.01% SDS, 1.1% Triton X-100, 1.2 mM EDTA, freshly complemented with 1x cOmplete^TM^ EDTA-free Protease Inhibitor Cocktail (Roche)). Following a 30 min preclear using blocked Protein A or G agarose beads (Roche), the chromatin was incubated with 5 µg antibody overnight at 4 °C (antibodies are listed in Table S6). Immune complexes were captured with 50 µl blocked Protein A or G agarose beads by incubation for 4 h with rotation and then washed with the following buffers: Low Salt Buffer (20 mM Tris pH8.1, 150 mM NaCl, 0.2% SDS, 1% Triton X-100, 5 mM EDTA, and 5% w/v sucrose), High Salt Buffer (50 mM HEPES pH 7.5, 500 mM NaCl, 0.1% deoxycholic acid, 1% Triton X-100 and 1 mM EDTA), Lithium Chloride Buffer (10 mM Tris pH 8.0, 250 mM LiCl, 0.5% deoxycholic acid, 0.5% NP-40 and 1 mM EDTA) and twice with TE Buffer (10 mM Tris-HCl pH 8.0, 1 mM EDTA). Captured chromatin was eluted in 125 µl elution buffer (1% SDS and 0.1 M NaHCO_3_) by shaking vigorously for 30 min at room temperature and then subjected to reverse crosslinking by incubating at 65 °C overnight in the presence of 200 mM NaCl. Input controls consisted of 50 µl (5%) samples of pre-cleared chromatin that were reverse crosslinked and purified as described for the IP samples. After incubation with 0.5 µg RNase A (1 h at 37 °C) and 20 µg Proteinase K (1 h at 45 °C), DNA was purified using the QIAquick PCR Purification kit (Qiagen).

For sequencing, ChIP DNA was further sonicated using a Bioruptor sonication system to obtain DNA fragments in the 100-300 bp range. Libraries were prepared and sequenced using the same procedure as described for the Native Chromatin Immunoprecipitation with exogenous spike-in experiments.

### RNA-seq sample preparation and sequencing

Total RNA was isolated from mES cells using the TRI Reagent (Sigma) following the protocol provided by the manufacturer. Contaminating genomic DNA was removed by incubating 10 µg RNA with 10 U DNaseI (Thermo Scientific) in Reaction Buffer for 30 min at 37 °C. The reaction was stopped by adding 10 µl of 50 mM EDTA after which RNA was extracted using a phenol-choloroform purification and ethanol precipitation. Material was quantified using RiboGreen (Invitrogen) on the FLUOstar OPTIMA plate reader (BMG Labtech) and the size profile and integrity analysed on the 2200 (Agilent, RNA ScreenTape). Input material was normalised to 1 µg prior to library preparation. Total RNA was depleted of ribosomal RNA using Ribo-Zero rRNA Removal Kit (Epicentre/Illumina, Human/Mouse/Rat) and library preparation was completed using NEBNext® Ultra Directional RNA Library Prep Kit for Illumina® (New England Biolabs), both following manufacturer’s instructions. Libraries were amplified for 12 cycles. Equivalent amounts of individual libraries were pooled together. Paired-end sequencing was performed using a Hiseq2000 51 bp platform (v3 SBS chemistry), generating a raw read count of >40 million reads per sample.

### Calibrated ChIP-seq computational analysis

ChIP-seq 75-bp paired-end reads were aligned to a concatenated mm10+galGal4 genome using bowtie2 (v2.2.5.0) with the options *--no-mixed* and *--no-discordant*. Reads that had multiple alignments were discarded and duplicate reads were removed. The number of reads mapping to the mm10 or galGal4 genome were counted and used to calculate the ratio of galGal4 to mm10 reads for both IP and inputs samples. To adjust for the variation in mouse cells used for the four experimental conditions from a single biological replicate, the number of galGal4 reads in the input samples were randomly down-sampled so that the resulting mm10:galGal4 ratio in the four input samples was equal (See Figure S1B and Table S1). Enriched regions were identified using the MACS2 function ‘callpeak’ (v2.0.10, Zhang et al., 2008) with the genome size set to 1.87 × 10^9^. Peak calling was performed independently for each biological replicate using all uniquely mapped mm10 reads. Regions identified as enriched in more than one sample from any condition were kept for downstream analyses.

To generate bigWig coverage files calibrated to the exogenous spike-in, we first randomly down-sampled BAM files containing the mm10 alignments for a particular sample based on the number of reads mapping to the galGal4 genome for that sample (in a manner which results in an equal mm10:galGal4 reads ratio in all samples). BigWig files were subsequently created using the Deeptools (Ramírez et al., 2016) function ‘bamCoverage’ (v2.4.2) with the options −*e* and −*bs 2*.

### ATAC-seq computational analysis

Whilst we incorporated spike-ins into our ATAC-seq experiments to facilitate calibration, the number of reads mapping to the exogenous galGal4 genome varied widely between the four biological replicates of a same experimental condition (Figure M1 below). Due to this variation and the lack of a consistent trend when comparing samples from different experimental conditions, we omitted the spike-in data from our analyses.

**Figure M1:**
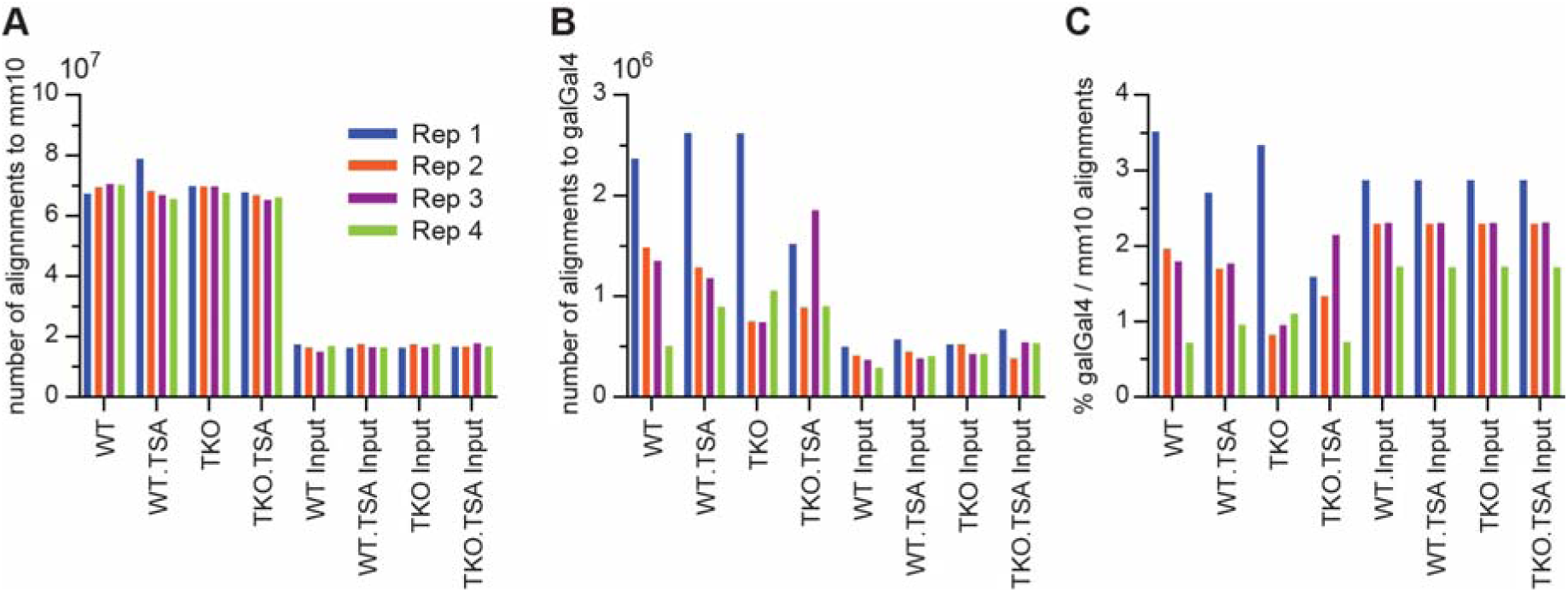
**A.** Number of uniquely mapped reads from each ATAC-seq library that align to the mm10 genome. **B.** Number of uniquely mapped reads from each ATAC-seq library that align to the galGal4 genome. **C.** Number of uniquely mapped reads from each ATAC-seq library that align to the galGal4 genome per 100 reads that align to the mm10 genome.

ATAC-seq 75 bp paired-end reads were trimmed using cutadapt (v1.16) to remove 3’ adapter sequences then aligned to a concatenated mm10+galGal4 genome using bowtie2 (v2.2.5.0) with the options *--no-mixed* and *--no-discordant*. Reads that aligned to the exogenous galGal4 genome, reads with multiple alignments and fragments mapping to a custom “blacklist” of artificially high regions of the genome were discarded. Duplicate reads were removed. Alignment files were randomly down-sampled to retain an equivalent number of alignments in all samples. BigWig coverage files were generated using the Deeptools (Ramírez et al., 2016) function ‘bamCoverage’ (v2.4.2) with the option −*bs* set to 2. Enriched regions were identified using the ‘dpeak’ peak-calling algorithm of DANPOS v2.2.2 (Chen et al., 2013) with the options −*m, –kd 250, -kw 150* and −*p 1e-60*. The R package DiffBind (Stark and Brown, 2011) was used to identify ATAC-seq peak regions with significant changes in enrichment between experimental conditions. Significant changes were determined using the DBA_DESEQ2 method applying a fold change cut-off of 1.5 and a p-value threshold of 0.01 after multiple-testing correction.

### Quantification of ATAC-seq reads within different categories of genomic regions

The genomic space was segmented into three categories of regions: “ATAC-seq peaks” that were called using DANPOS ‘dpeak’ as described above; “Repmask regions” which included all genomic intervals that were annotated as repetitive based on the UCSC RepeatMasker annotation but excluded sequences that overlap with ATAC-seq peaks; and a “Unannotated” category which grouped together all remaining genomic intervals. The bedtools “intersect” command was used to count the number of alignments from each sequencing dataset that overlapped regions within each category. All alignments to the mm10 genome were counted, including those marked as having multiple alignments. Reads that overlapped two regions were counted only once: reads that only partially overlapped with an “ATAC peaks” region were included in this category while reads that overlapped a “Repmask” region and an “Unannotated” region were counted for the “Repmask” region only. We then calculated the percentage of total reads in the dataset that mapped to each category (Figure 2E).

To estimate the percentage of reads mapping outside of “ATAC peak” regions that would align to the “Repmask” and “Unnanotated” regions by chance (random or expected distribution), we shuffled the reads with no alignment to ATAC peaks within the combined “Repmask” and “Unnanotated” genomic space (using the bedtools “shuffle” command) and repeated the counting procedure (Figure S2E). We calculated “observed/expected” values based on the number of reads mapping to these two categories before and after shuffling (Figure 2F).

### ChIP-seq computational analysis

ChIP-seq 75-bp paired-end reads were aligned to the mm10 genome using bowtie2 (v2.2.5.0) with the options *--no-mixed* and *--no-discordant*. Reads that had multiple alignments were discarded, duplicate reads were removed as well as fragments mapping to a custom “blacklist” of artificially high regions of the genome (derived from the ENCODE Consortium Project, 2012). All alignment files from the same transcription factor ChIP were randomly down-sampled to match the number of alignments in the minimal ChIP or input sample. BigWig coverage files were generated using the Deeptools (Ramírez et al., 2016) function ‘bamCoverage’ (v2.4.2) with the options −*e* and −*bs 2*. Enriched regions were identified using the ‘dpeak’ peak-calling algorithm of DANPOS v2.2.2 (Chen et al., 2013) with the options −*m, -kd 250, -kw 150* and −*p 1e-100*.

When visualising the GABPA ChIP-seq datasets we noticed that sequencing reads were not only enriched in a localised manner, forming the narrow peaks typically seen in transcription factor ChIP-seq data, but also within broader domains whose boundaries coincided closely with the edges of transcribed genes, particularly those with high levels of transcription. For the purposes of this study, we restricted our analysis to localised sites of GABPA ChIP-seq enrichment. While the ‘dpeak’ peak-calling algorithm is designed to identify shorter enriched regions rather than larger domains, many of the focal “peaks” called using the parameters noted above were located within the broader GABPA enriched regions. In order to filter these out, we first identified genomic domains >2000bp broadly enriched with GABPA using the DANPOS function ‘dregion’ with the options *-m 1 -rd 1500 -rw 2000* and *-p 1e-50*. Subsequently, peaks called using dpeak (n = 10022) that overlapped with a called region were discarded unless they intersected a gene promoter (−1kb to 100bp around TSS). 4735 peaks remained, 74.7% contained a GABPA motif, compared to 56.9% before filtering.

The R package DiffBind (Stark and Brown, 2011) was used to identify ChIP-seq peak regions with significant changes in enrichment between experimental conditions. Significant changes were determined using the DBA_DESEQ2 method applying a fold change cut-off of 4 and a p-value threshold of 10^−3^ after multiple-testing correction (Bonferroni correction).

### RNA-seq computational analysis

RNA-seq 51 bp paired-end reads were aligned to the mouse mm10 genome using TopHat (v2.0.13) (Trapnell et al., 2009) with the *--library-type* option *fr-firststrand*, *--mate-inner-dist* set to *100* and coverage based searching for junctions disabled. FeatureCounts v1.4.5-p1 (Liao et al., 2014) was used to count aligned RNA-seq reads that mapped to the exons of annotated genes. Read counting was strand specific (option *-s* set to 2). Reads with a mapping quality below 1 were excluded. Differential analysis was performed using the R package edgeR (v3.22.3, McCarthy et al., 2012; Robinson et al., 2010) with raw read counts supplied as input. Genes with a low level of expression in all samples were filtered out using a counts per million (cpm) threshold of 1. The generalized linear model (GLM) likelihood test was used to determine differential expression between multiple groups of biological replicate samples. Genes or transcripts were considered as significantly differentially expressed if they had a p-value < 10^−5^ after adjusting for multiple comparisons using the Benjamini and Hochberg procedure. Strand-specific bigWig coverage files were generated using the Deeptools (Ramírez et al., 2016) function ‘bamCoverage’ (v2.4.2) with the options −*-normalizeUsingRPKM, --filterRNAstrand* and −*bs 1*.

The enrichment analysis of genes with tissue specificity was performed using the DAVID functional annotation tool (version 6.8, Huang et al., 2009a, 2009b). A functional annotation chart based on the Uniprot UP_TISSUE annotation was reported using default options.

### Quantification of reads mapping to repetitive genomic features and differential analysis

For individual ATAC-seq, ChIP-seq or RNA-seq sample, total read counts for every type of repetitive element were generated using a custom script, as detailed below. All bowtie2 alignments, including those ascribed to multi-mapping reads, were taken into account. The position and annotation of interspersed repeats and low complexity DNA sequences in the mm10 genome was obtained from UCSC (created using RepeatMasker, from the RepBase library of repetitive elements (Jurka et al., 2005)). The custom script makes use of the Samtools “view” command to count reads at genomic intervals, the Bedtools “nuc” command to determine the CpG content of genomic intervals and the ‘st’ tool (https://github.com/nferraz/st) to calculate CpG content and nucleotide substitution statistics across all copies of a repeat element type. The R package DESeq2 v1.20.0 (Love et al., 2014) was used to identify repeat element types that show significant differences in coverage between conditions. A p-value threshold of < 0.05 was used to identify significant differences after correction for multiple testing using the Benjamini & Hochberg procedure.

### Misc

Metaplots used to display and summarise ATAC-seq, ChIP-seq or Native H3ac ChIP-seq signals were generated using Deeptools functions (Ramírez et al., 2016). As input, these tools were provided with genome coverage files (bigWigs) created after merging alignments from biological replicate samples.

The gene annotation files for mm10 or galGal4 used in this study were downloaded from UCSC. Genomic regions, for example ChIP-seq or ATAC-seq peaks, were annotated using the HOMER script annotatePeaks.pl (Heinz et al., 2010).

The HOMER script findMotifsGenome.pl (v4.7, Heinz et al., 2010) was used to define *de novo* motifs that were enriched within particular sets of target genomic intervals. The same script was used to screen for the enrichment of previously known or custom motifs. The maximum log-odds score for a given position weight matrix (PWM) motif within each interval in a .BED file was determined using the HOMER annotatePeaks.pl script with the *-mscore* option.

All boxplots were generated using the R package ggplot2. The median is indicated by a bar, the lower and upper hinges correspond to the first and third quartiles while the upper/lower whisker extend from the hinges to the largest value no further than 1.5x the inter-quartile range.

### Analysis of publicly available WGBS data

Whole genome bisulfite sequencing data from E14 mouse ESCs (GSM1027571, Habibi et al., 2013) was analysed using Bismark (v 0.12.5). Only cytosines with a read coverage of five or more were taken into account in this study.

### Data and Code Availability

ChIP-seq, ATAC-seq and RNA-seq sequencing data generated during this study are available on Gene Expression Omnibus (accession GSE131366).

## Supplemental Information

**Figure S1:**
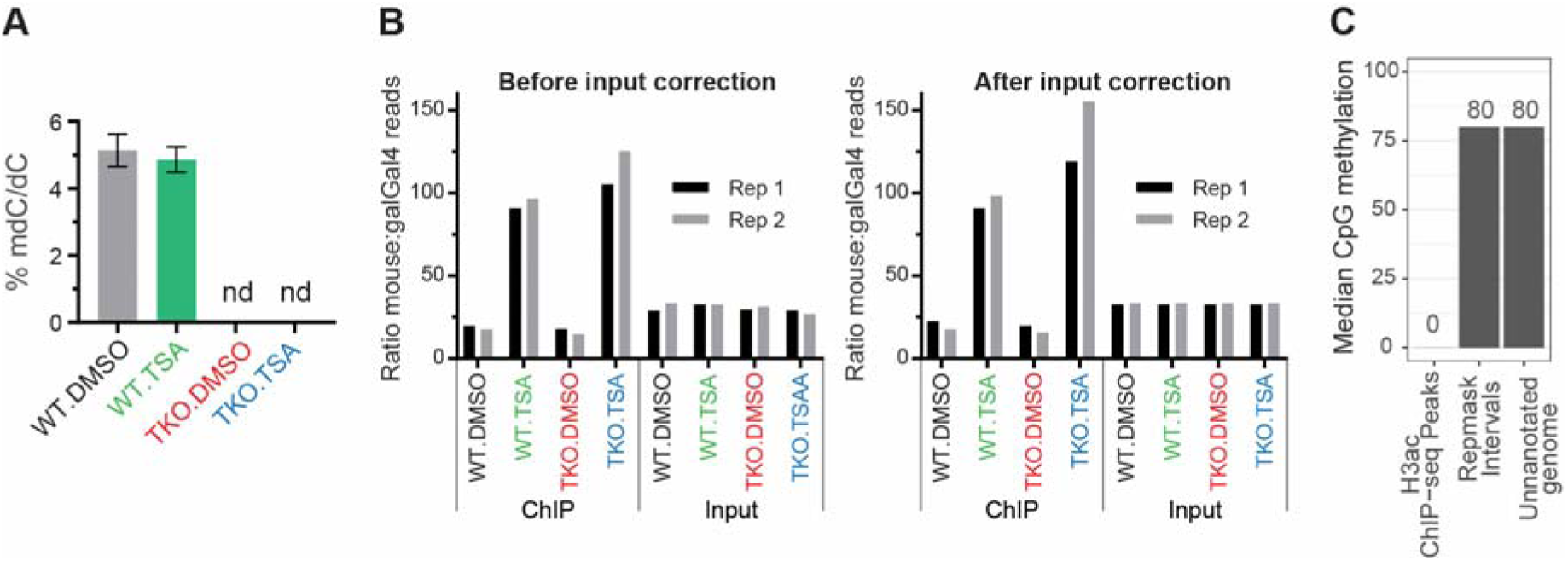
Disruption of HDAC activity and DNA methylation in mESCs. Related to Figure 1. **A.** Mass spectroscopy measurements of 5mC in DMSO or TSA-treated wild-type and DNMT.TKO cells shown as a percentage of total cytosine. Data are represented as mean ± SD. nd: not detected. **B.** Number of reads mapping to the mm10 versus galGal4 genomes for each sample. Data are shown before and after adjusting for the variation within input samples (see Materials and Methods and Table S1). **C.** Median CpG methylation levels for genomic intervals separated into three mutually exclusive categories. Whole genome bisulfite sequencing data from E14 mESCs was obtained from Habibi et al., 2013. Exact values are shown above the bars.

**Figure S2:**
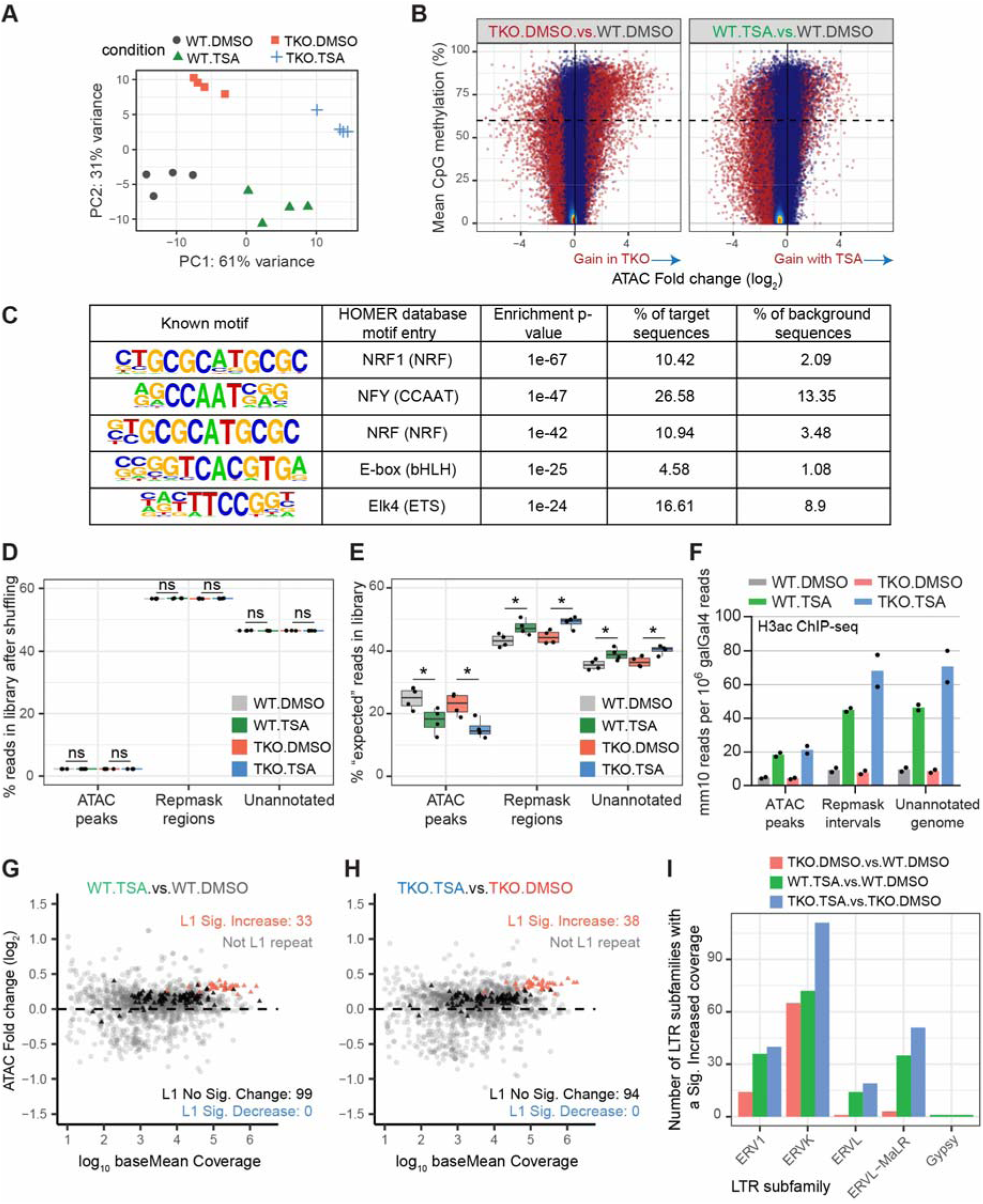
Distinct contributions of DNA methylation and HDAC activity to the chromatin accessibility landscape. Related to Figure 2. **A.** Principle component analysis (PCA) plot generated using regularised log-transformed read counts from all ATAC-seq samples at every THS. The R package *DESeq2* (Love et al., 2014) was used for plotting. **B.** Scatterplots comparing the change in ATAC-seq signal with the average CpG methylation levels at 83395 ATAC-seq peak intervals. Regions with significantly differential accessibility (fold change > 1.5 and FDR < 0.01) are shown in red. The dashed line indicates 60% CpG methylation. **C.** The known motifs most enriched in DNMT.TKO specific ATAC-seq peaks (target sequences) relative to background sequences. The HOMER motif database entry provides information about the transcription factor’s DNA-binding domain (DBD-family in brackets). Target sequences were restricted to those that were methylated (≥ 60% mean CpG methylation) and had at least a four-fold increase in accessibility compared to WT cells. See also Table S5. **D.** Boxplots summarising the percentage of reads from each ATAC-seq library that map to intervals split into three mutually exclusive categories, after having randomly shuffled the reads throughout the genome. Two-tailed Student t-tests were used to determine significant differences: *ns* = non-significant (p-value > 0.05). **E.** Boxplots summarising the percentage of reads from each ATAC-seq library that map to intervals split into three mutually exclusive categories, after having randomly shuffled the non-peak reads within the “Repmask” and “Unannotated” regions. Two-tailed Student t-tests were used to determine significant differences: * p-value < 0.05; ** p-value < 0.01; *ns* = non-significant (p-value > 0.05). **F.** Distribution of H3ac ChIP-seq reads within genomic intervals separated into three mutually exclusive categories. For every sample, the number of mm10 reads overlapping each category was divided by the total number of reads mapping to the galGal4 genome. Data are represented as mean; points indicate the values for two biological replicates. **G.** MA plots comparing for each type of repetitive element, the fold change in ATAC-seq signal across all genomic copies to the baseMean ATAC-seq signal. LINE-1 elements are highlighted. ATAC-seq samples generated from TSA- or DMSO-treated wild-type mESCs were compared. Differential analysis was performed using the R package DESeq2 (Love et al., 2014). Changes were deemed significant with an FDR < 0.05). See also Table S3. **H.** Same as (G), except samples from TSA- and DMSO-treated DNMT.TKO mESCs were compared. **I.** Number of LTR element types, subdivided by family, that show significantly increased ATAC-seq signal for three pairwise comparisons. See also Table S3.

**Figure S3:**
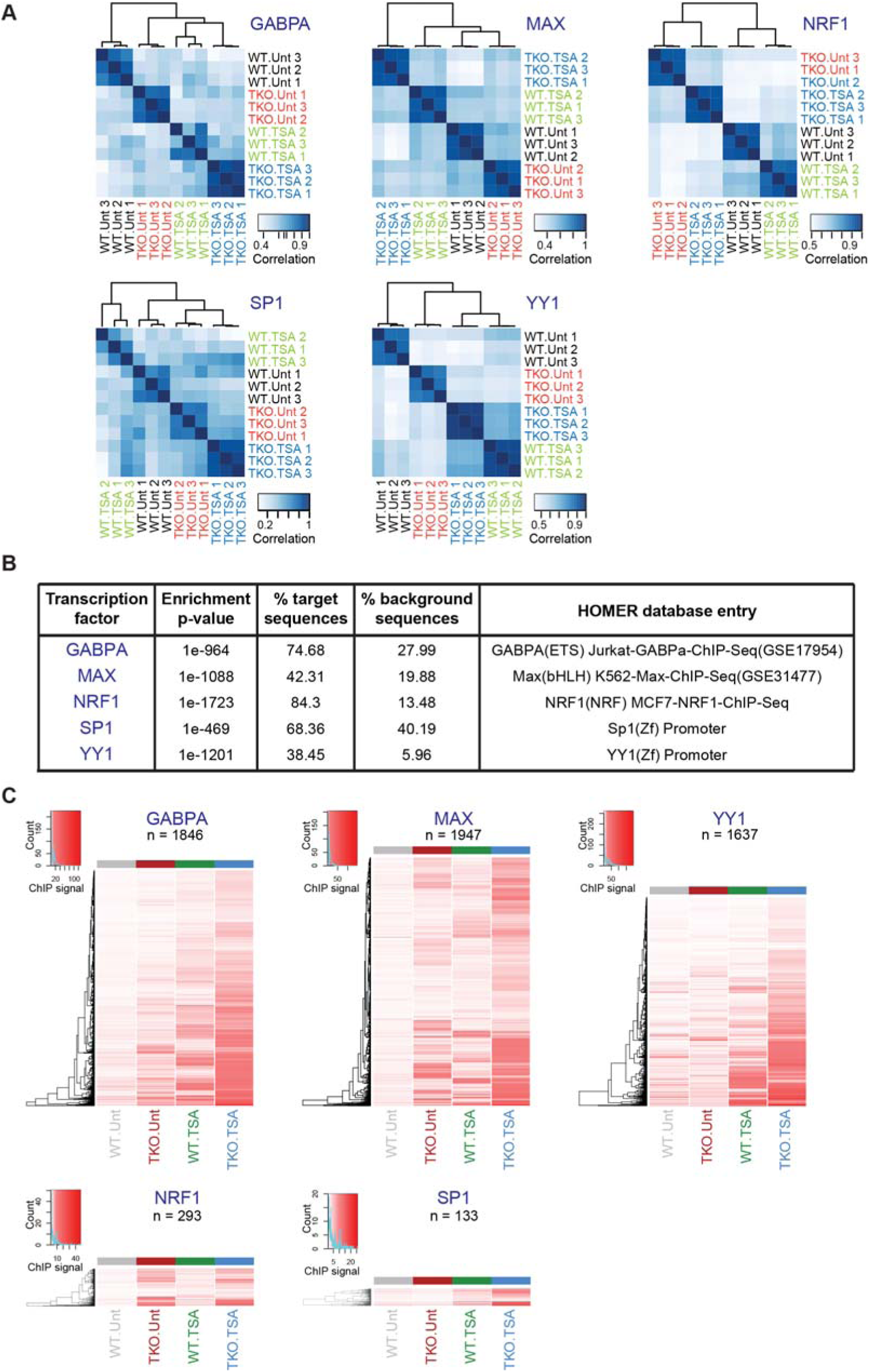
DNA methylation and HDAC activity can modulate transcription factors occupancy. Related to Figure 3. **A.** Correlation heatmaps comparing the ChIP-seq signal obtained for every sample at all identified binding sites. ChIP signal was calculated as the RPKM of the TF ChIP-seq divided by the RPKM of the input control. Pearson correlation coefficients were calculated for each pair of samples and these values were used for clustering and the colour scale. Correlation heatmaps were generated in R using the *dba.plotHeatmap()* function of the *DiffBind* package (Stark and Brown, 2011). **B.** Most enriched motifs found within transcription factors ChIP-seq peaks (target sequences) relative to background sequences. For all factors except GABPA, the motif shown is the most enriched compared to all others entries in the HOMER collection. This GABPA motif is ranked 5th most enriched, however the top 10 motifs enriched at GABPA peaks are all highly similar ETS factor motif (See Table S5). **C.** Heatmaps show the ChIP-seq signal for every peak region at which a significant difference in occupancy was detected between two dataset pairs (TKO.Unt vs. J1.Unt; J1.TSA vs. J1.Unt; TKO.TSA vs. TKO.Unt; or TKO.TSA vs. J1.Unt). ChIP-seq signal was calculated for each sample as the RPKM of the TF ChIP-seq divided by the RPKM of the input control. Values from three biological replicates were averaged for plotting. The heights of the heatmaps are scaled to the number of peak intervals.

**Figure S4:**
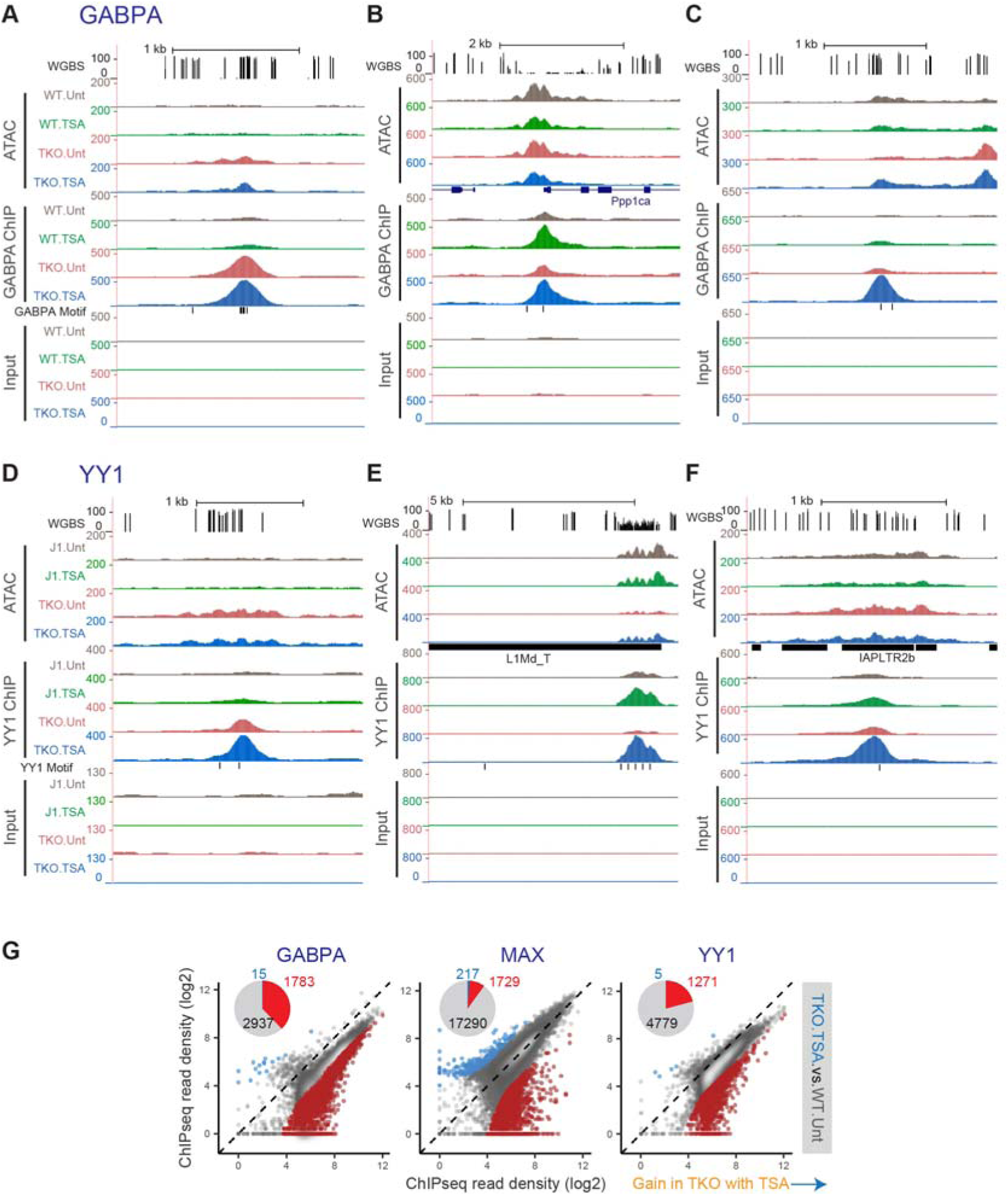
DNA methylation and HDAC activity can modulate transcription factors occupancy. Related to Figure 3. **A.** Representative UCSC genome browser snapshot showing CpG methylation levels (Habibi et al., 2013), ATAC-seq, GABPA ChIP-seq and Input ChIP-seq read coverage. The position of genes and that of sequences that match the GABPA motif are shown. ATAC-seq and ChIP-seq coverage graphs were generated after merging alignments from replicate samples. Mm10 coordinates: chr1:185,055,705-185,059,261. **B.** Same as (A). Mm10 coordinates: chr19:4,188,845-4,196,031. **C.** Same as (A). Mm10 coordinates: chr8:24,336,044-24,339,393. **D.** Representative UCSC genome browser snapshot showing CpG methylation levels (Habibi et al., 2013), ATAC-seq, YY1 ChIP-seq and Input ChIP-seq read coverage. The position of repeat elements and that of sequences that match the YY1 motif are shown. ATAC-seq and ChIP-seq coverage graphs were generated after merging alignments from replicate samples. Mm10 coordinates: chr11:74,950,700-74,954,985. **E.** Same as (D). Mm10 coordinates: chr4:49,457,807-49,470,813. **F.** Same as (D). Mm10 coordinates: chr6:34,330,141-34,333,726. **G.** For every occupancy peak identified for GABPA, MAX or YY1, normalised ChIP-seq signal was plotted for samples generated from TSA-treated DNMT.TKO cells (TKO.TSA) versus untreated wild-type cells (WT.Unt). Regions with significantly differential occupancy (fold change > 4 and adjusted p-value < 10^−3^) are coloured on the scatter plots and their numbers are summarised in the form of pie charts (light blue = significant decrease; red = significant increase).

**Figure S5:**
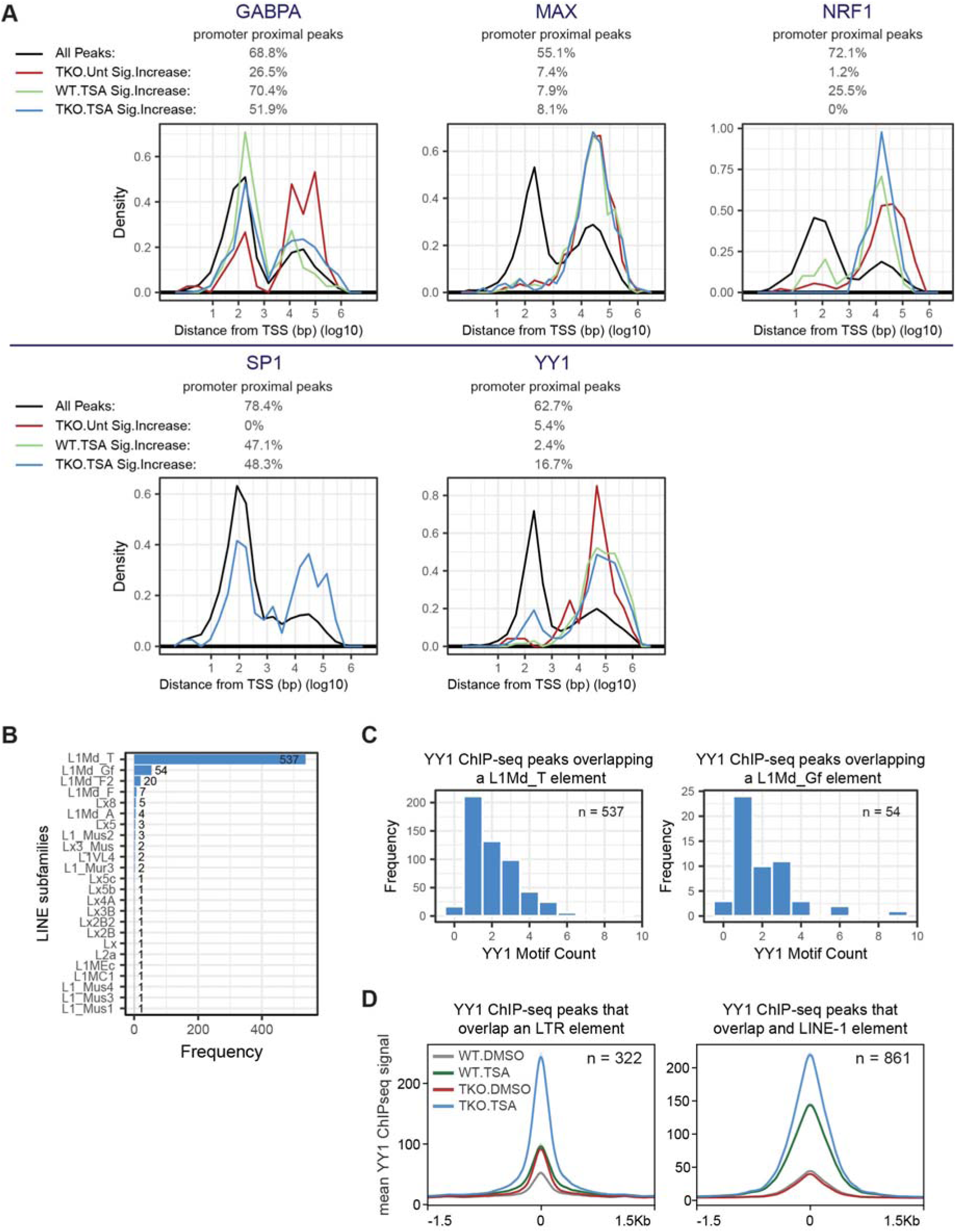
Characteristics of transcription factor binding sites. Related to Figure 4. **A.** Density plots showing the distance of ChIP-seq peaks to the nearest transcription start site (TSS). The percentages of peaks that are proximal to the TSS (centre of peak within 1500 bp of TSS) are indicated above the plots. **B.** The extent of overlap between TKO.TSA-specific YY1 ChIP-seq peaks and different subtypes of LINE elements. Only peaks with significantly increased YY1 occupancy in TSA-treated versus untreated DNMT.TKO cells and that overlap a LINE are included (n = 652). **C.** Number of TKO.TSA-specific YY1 ChIP-seq peaks that harbour a cognate binding motif. Left: peaks that overlap an L1Md_T element only. Right: peaks that overlap an L1Md_Gf element only. **D.** Metaplots showing the average YY1 ChIP-seq signal in 3 kb regions surrounding the centre of ChIP-seq peaks that overlap either an LTR (left) or LINE (right) elements.

**Figure S6:**
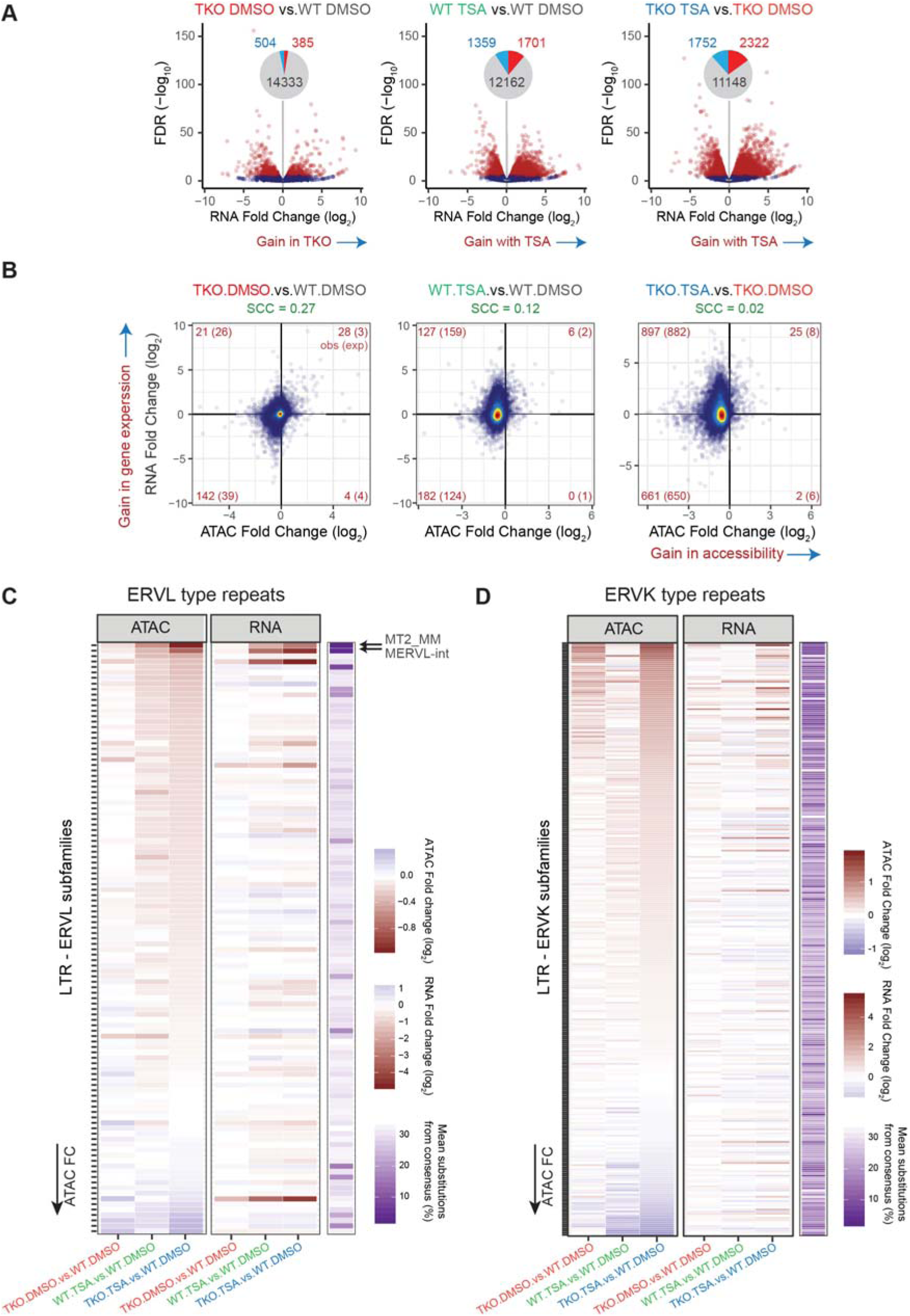
The impact of DNA methylation loss and HDAC inhibition on gene expression. Related to Figure 6. **A.** Volcano plots representing the false discovery rate (FDR) and fold change values obtained through pairwise differential analyses of strand-specific RNA-seq read counts at 15222 genes. Genes with significantly differential expression (adjusted p-value < 10^−5^) are shown in red on the scatter plots and their numbers are summarised in the form of pie charts (light blue = significant decrease; red = significant increase). **B.** Changes in accessibility at promoter-proximal THS regions (centre of peak within 1500 bp of TSS) were compared to changes in expression at their closest gene. The analysis was performed on 13068 THS:gene pairs involving 12356 genes. The numbers of ATAC-seq peaks associated with significant changes in both accessibility <u>and</u> gene expression are indicated in red in each quadrant. In brackets are indicated the expected number of sites showing both significant changes in accessibility and gene expression based on the total number of significant differential events. SCC = Spearman Correlation Coefficient. **C.** For each ERVL subtype (N = 109), we plotted the fold change in ATAC-seq or RNA-seq signal along with scores relating to their sequence conservation (purple). ERVL subtypes were sorted based on the fold change in ATAC-seq coverage when comparing DNMT.TKO-TSA to wild-type cells. See also Table S3. **D.** For each ERVK subtype (N = 231), we plotted the fold change in ATAC-seq or RNA-seq signal along with scores relating to their sequence conservation (purple). ERVK subtypes were sorted based on the fold change in ATAC-seq coverage when comparing DNMT.TKO-TSA to wild-type cells. See also Table S3.

**Table S1:** Number of raw and mapped reads for all high-throughput sequencing samples.

**Table S2:** Lists of enriched regions for ATAC-seq and ChIP-seq experiments along with differential analyses results and annotations.

**Table S3:** Raw read counts and differential analyses results at repetitive elements.

**Table S4:** Results of differential gene expression analyses.

**Table S5:** Complete results from motif enrichment and functional annotation analyses.

**Table S6:** List of antibodies used in ChIP-seq experiments. Nucleoside transition and retention times for Mass Spectrometry analysis.

